# Sleep-like behavior is a fundamental property of the tripartite synapse

**DOI:** 10.1101/2020.02.02.917633

**Authors:** Shubhada N Joshi, Aditya N Joshi, Narendra D Joshi

## Abstract

The tripartite synapse, consisting of the presynaptic neuron, post-synaptic neuron, and an astrocyte, is considered to be the main locus of signaling between neurons in the brain.^1,2^ Neurotransmission is energetically very expensive^3,4^, and the primary neurotransmitter utilized for signaling is glutamate. It has been found that glutamate is also used as a substrate for energy generation.^5,6^ However, it is unclear what the relationship is between energy generation and availability of neurotransmitter during glutamatergic neurotransmission. Here we show that availability of energy, represented by adenosine triphosphate (ATP), and glutamate for neurotransmission are intimately related, and in fact determine the ability to signal at the tripartite synapse. Using a novel neurochemical mathematical model of the tripartite synapse, we found that glutamate concentrations for neurotransmission and ATP concentrations were interdependent, and their interplay controlled the firing pattern of the presynaptic terminal, as defined by synaptic vesicle release. Furthermore, we found that depending on the parameters chosen in the model, the tripartite synapse demonstrated behavior with limit cycles, alternating between high- and low-frequency firing rates. Our results show that complex behavior with high- and low-activity states, qualitatively meeting the characteristics of sleep^7^ emerges directly from the nature of the tripartite synapse, with glutamate and ATP concentrations serving as the signals for state changes. We anticipate that our model will serve as a starting point to further elucidate the energetics of neuronal and brain functioning, and eventually shed light on the fundamental question of the nature and necessity of sleep.

## Main

The mammalian brain is an information processing unit, and neurons are considered the workhorses of this function. There are billions of neurons in the mammalian brain, but there are at least as many, and possibly more, glia, including astrocytes.^4,8^ In more recent years, the crucial role of astrocytes in neuronal synaptic transmission has been elucidated, giving rise to the concept of the “tripartite synapse”.^1,2,9^

Approximately 80% of neocortical neurons use glutamate as their neurotransmitter, and approximately 85% of neocortical synapses are excitatory. The supply of glutamate for neurotransmission is thus essential for the functioning of the brain.^5,6^ For a signal to be processed at a glutamatergic synapse, previously-released glutamate must be removed efficiently from the synaptic cleft, and neurons must replenish their supplies of glutamate. Astrocytes, as participants in the tripartite synapse, are intimately involved in the recycling of glutamate through the glutamate-glutamine cycle: the released glutamate is mostly reuptaken by astrocytes, converted into glutamine, exported back to neurons, and then converted back to glutamate for further neurotransmission.^6^

The glutamate-glutamine cycle was shown to be highly active in cerebral cortex in a rat model, and it was demonstrated that incremental glucose oxidation was tightly linked to this cycle in an approximately 1:1 stoichiometry. In the resting human brain, the cycle constitutes a major metabolic flux, accounting for approximately 80% of glucose oxidation.^10^ Neurons are themselves also very metabolically active due to significant energy expended for maintaining and resetting the resting membrane potential for signaling.^3,4^ Glutamatergic signaling thus represents a major energy expenditure for both neurons and astrocytes. The universal energy currency for all cells is adenosine triphosphate (ATP). In brain tissue, ATP is produced locally through glycolysis, oxidative metabolism using acetyl-Coenzyme-A (acetyl CoA) in the mitochondria, or through non-local transport of lactate produced via glycolysis in the astrocytes. This lactate is subsequently oxidized in the presynaptic glutamatergic terminals.^11,12^ Notably, however, in addition to being the major excitatory neurotransmitter in the brain, glutamate is also used as a substrate for production of ATP.^6,10^ This allows it to be a functional linkage between neurotransmission and energy availability in the tripartite synapse.

We developed a novel parametric model of astrocyte-neuron-vasculature-extracellular-fluid derived from first principles of intracellular chemical processes, to demonstrate the intimate connection between energetics and neurotransmission in the tripartite synapse. In this work, we demonstrate the appearance of several unique properties of a neuronal system, including use-dependence of the frequency of firing at the synapse which depends on both ATP concentration and availability of neurotransmitter, and the presence of “UP” (high firing frequency) and “DOWN” (low firing frequency) states of neuronal firing.^13^ We also find that the system is “chaotic”—small changes in a single parameter can have long-range implications, and there is an emergence of “limit cycles” which describe oscillatory behavior of the system. With this work we lay the groundwork for our overarching hypothesis, that oscillations between rest and activity are an emergent property of the neuronal-astrocyte system. Ultimately, this model can be expanded to encompass larger units within the brain, and may demonstrate that oscillatory periods of rest and activity (i.e. *sleep and wakefulness*) arise as fundamental properties of complex neuronal networks. We further discuss the motivations for studying sleep, examine several other prior models, introduce the topic of diffuse modulatory systems in sleep, and comment on potential implications, in the Supplementary Discussion.

## Model Description

We modeled a presynaptic terminal, synaptic cleft, and astrocytic process of the “tripartite synapse”; the post-synaptic terminal was not considered as it is more relevant to signaling than to energetics of the presynaptic neuron and astrocyte assembly. There are five compartments in this model: presynaptic terminal (of the presynaptic neuron), astrocytic process, synaptic cleft, extracellular fluid (ECF), and blood vessel. A schematic of the model is shown in Figure 1. In the presynaptic terminal, there were three primary processes modeled: reactions related to energy substrate, reactions related to neurotransmission, and reactions linking energy utilization with neurotransmission. Processes related to energy in the presynaptic terminal included uptake of lactate from the ECF, the reactions of the tricarboxylic acid (TCA) cycle, production of ATP from the electron transport chain, use of ATP for “vegetative” or “housekeeping” functions (such as transcription, protein synthesis, and other processes not directly related to neurotransmission^14^), and diversion of energy substrate intermediates for biosynthetic processes (such as amino acid synthesis). In keeping with the astrocyte-neuron lactate shuttle (ANLS) hypothesis^15^, we did not include glycolysis in the presynaptic terminal (see Methods). Neurotransmitter-related reactions in the presynaptic terminal included glutamine uptake from the ECF, deamination of glutamine to glutamate, vesicle loading, vesicular release, and “vegetative” use of glutamine and glutamate (such as protein synthesis^6^). Processes linking energetics with neurotransmission in the presynaptic terminal included the interconversion of α-ketoglutarate and glutamate, and the function of the sodium-potassium pump to restore the resting membrane potential, which utilizes ATP. The membrane potential was calculated based solely on the charge distribution created by movement of the sodium and potassium ions across the presynaptic terminal cell membrane.

**Figure 1.**
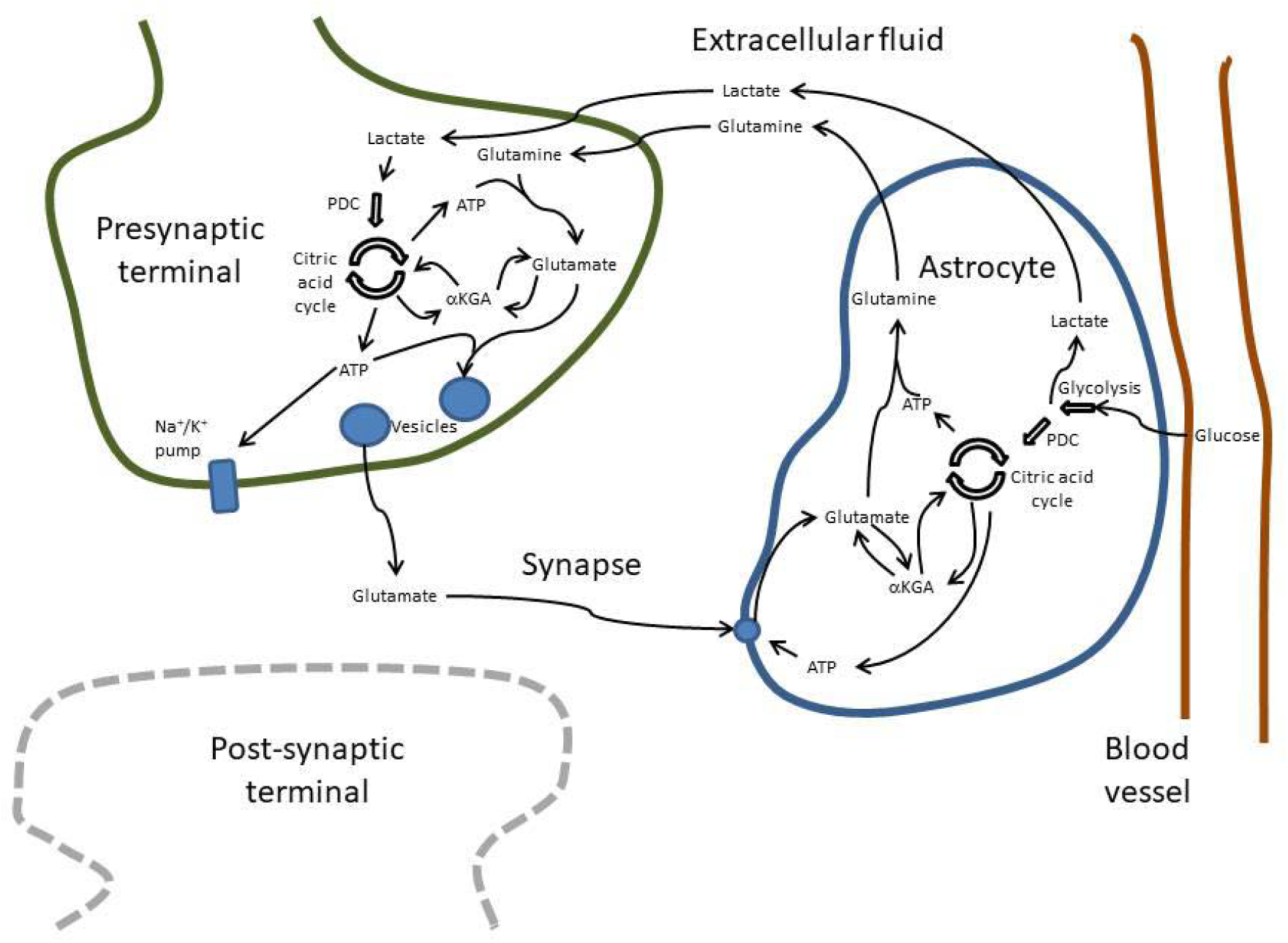
Schematic of the model. The presynaptic terminal, astrocyte, extracellular fluid, and vascular compartments are modeled; the post-synaptic terminal is depicted but is not part of the model. The major energy-generating processes and processes important for neurotransmission are shown, as well as the reactions that link the two. The heavy arrows represent well-described sequences of metabolic reactions. PDC—pyruvate dehydrogenase complex; ATP—adenosine triphosphate; *α*KGA—*α*-ketoglutarate; Na^+^/K^+^ pump—ATP-dependent sodium-potassium exchanger pump. This tripartite synapse is part of a larger network of similar synapses, whose connectivity properties are included in the parameter *ζ*.

In the astrocytic compartment, there were similarly three primary processes modeled: energy reactions, neurotransmission reactions, and reactions linking the two. Energy reactions included uptake of glucose from the blood vessel, glycolysis, the TCA cycle, the electron transport chain, the anaplerotic synthesis of oxaloacetate from pyruvate (catalyzed by pyruvate carboxylase^16^), export of lactate into the ECF (ANLS)^15^, “vegetative” use of ATP, and diversion of energy substrate intermediates for biosynthetic processes. Neurotransmitter-related reactions included uptake of glutamate from the synapse, amination of glutamate to glutamine, glutamine export to the ECF, and “vegetative” use of glutamine and glutamate.^6^ The primary reaction linking energetics to neurotransmission in the astrocyte is the interconversion of α-ketoglutarate and glutamate.

The processes modeled in the remaining three compartments (synaptic cleft, ECF, and blood vessel) were transport processes. In the synaptic cleft, the following processes occurred: release of vesicular glutamate into the synapse, glutamate reuptake from the synapse into the astrocyte, and diffusion of glutamate away from the synapse. In the ECF, there was glutamine release from astrocytes and uptake into neurons as described in the glutamate-glutamine cycle^6^, as well as losses of glutamine to diffusion. The ECF also included lactate release from the astrocytes and uptake by neurons per the ANLS hypothesis.^15^ The ECF served as the compartment from which sodium and potassium ion fluxes occurred during neuronal signal transmission. The blood vessel served as the compartment from which the astrocyte imported glucose.

Signal processing was modeled solely in the presynaptic terminal. Specifically, the arrival of synaptic inputs at the dendrites and soma, summation at the axon hillock, and axonal transmission of action potentials was abstracted into a single summary input frequency. That is, the “input frequency” represented the average frequency of action potentials that would arise as a result of the summation of signals at all of the dendritic and somatic synapses. The input frequency was comprised of two components: a constant component (*ε*) and a variable component (*ν*). The constant component represented a spontaneous firing frequency, representing the low rate of synaptic vesicle release that occurs spontaneously in neurons.^17^ The variable component of the input frequency represented the contribution of the network to which the presynaptic terminal belongs. We define a value *φ* as the ratio of the moving average output frequency (calculated as the average frequency of *synaptic vesicle release*, a surrogate for successfully transmitted action potentials, over the past three firing events) to the combined maximum of the input frequency (*ε* + *ν*). The actual input frequency takes one of two values depending on *φ* and another critical value *ζ*, which represents the fraction of other synapses in the network that need to have sufficient energy and neurotransmitter resources in order for the entire network to continue running at a high frequency. If *φ* decreases to a value less than *ζ* because of a lack of either energy or neurotransmitter resources, this is taken to be a surrogate for the condition that other synapses in the network also do not have enough resources to sustain a high firing frequency. In this case, the network contribution to the input frequency becomes negligible and input frequency equals *ε*. If *φ* is greater than *ζ*, then the entire network is able to continue running at a higher frequency, allowing the total input frequency to remain (*ε* + *ν*). If the network contribution is negligible (input frequency is *ε*), but the presynaptic terminal has sufficient resources in terms of ATP and vesicles, this is taken as a surrogate that enough other synapses in the network have the resources to “restart” the network, causing the input frequency to become (*ε* + *ν*) again as an “ignition” phenomenon. A further discussion of this “ignition” phenomenon in our model is found in the Supplementary Discussion.

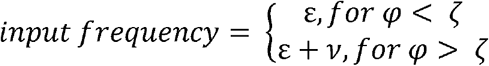

The “firing signal”—the signal that indicates that the presynaptic terminal should release a synaptic vesicle—was calculated based on the input frequency. Specifically, a synaptic vesicle would be released if the modulus of the simulation time with respect to the inverse of the input frequency was within a certain window. That window represented the time constant after which an action potential reaching the presynaptic terminal would dissipate without causing a vesicular release (a time constant we have called *χ*). Window effects are further explored in the Supplementary Discussion. In addition to being within the time window to fire, there would also have to be sufficient ATP and a sufficient number of vesicles for the synaptic vesicle to be released.

## Rest and Activity

We simulated the behavior of the presynaptic terminal-astrocyte assembly at low and high values of the input firing frequency. In a representative simulation of a lower input frequency (Figure 2), it can be seen that the behavior of the system does not change during the entire duration of the simulation. Figure 2a depicts that neither presynaptic ATP nor the number of vesicles is significantly depleted (middle and top traces respectively; dashed line indicates ATP threshold for firing in middle trace), while membrane potential remains at a single baseline during the simulation, punctuated by regular increases at the time that the neuron releases a synaptic vesicle—indicating the arrival of action potentials (bottom trace, and inset). In Figure 2b, the moving average output frequency (top trace) is seen to be identical to the input frequency (bottom trace).

**Figure 2.**
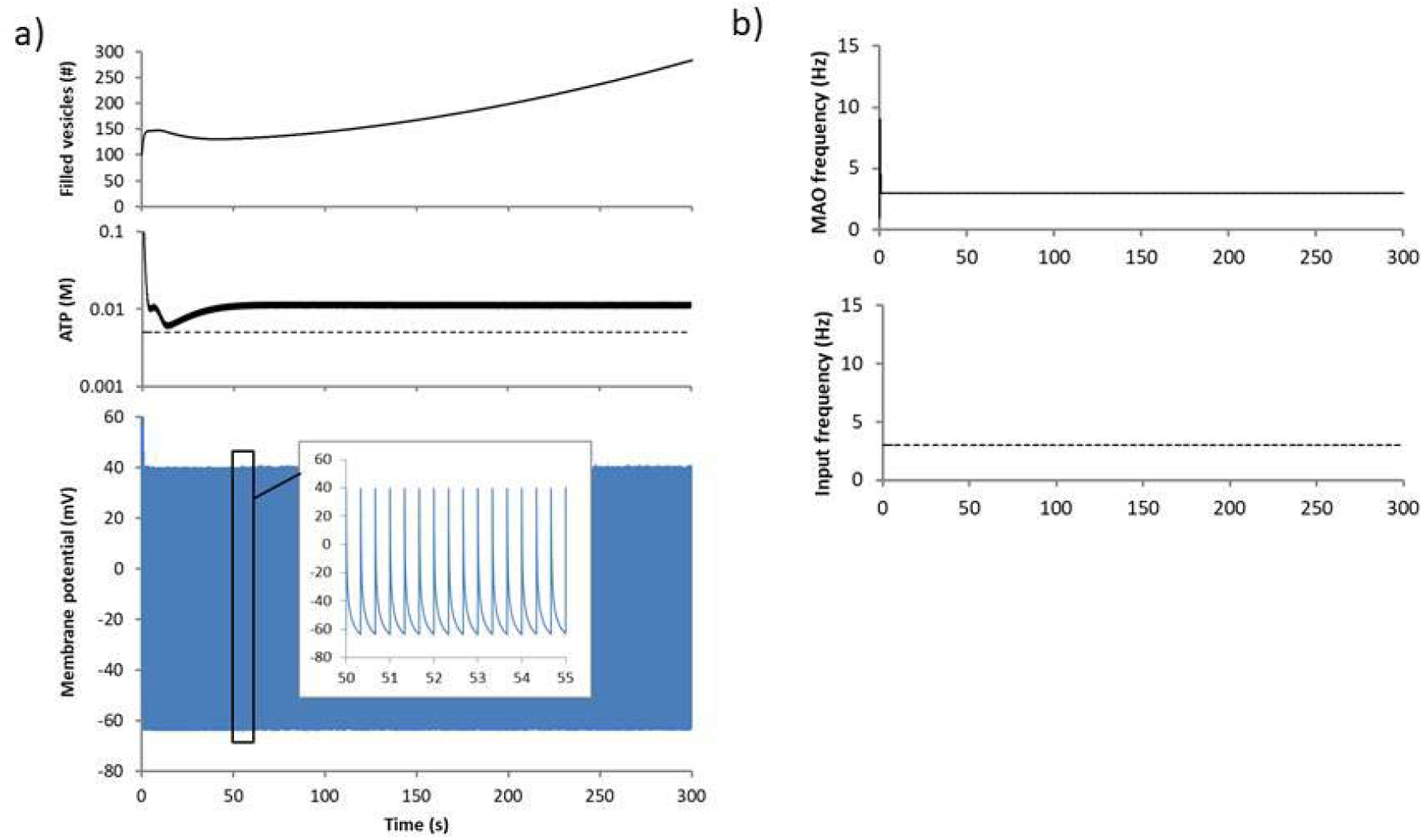
Input-limited operation. Regime A. *ε* = 1 Hz, *ν* = 2 Hz, *ζ* = 0.6, *χ* = 25 ms. Values shown are for the presynaptic terminal. a) *Top trace*: filled vesicles as a function of time. *Middle trace*: ATP concentration as a function of time. *Bottom trace*: membrane potential as a function of time. *Inset*: zoomed in view of membrane potential as a function of time. b) *Top trace*: moving average output frequency (“MAO frequency”) as a function of time. *Bottom trace*: input frequency as a function of time.

The behavior of the system significantly changes with a higher input frequency (Figure 3); “lower” and “higher” frequencies are relative, and depend on the specific parameters chosen (*data not shown*). As is seen in Figure 3a, the membrane potential now demonstrates *two* baseline levels, a higher (less negative) potential during active periods (“UP” states), and a lower (more negative) potential during quiescent periods (“DOWN” states), which initially alternate rapidly (bottom trace). After approximately 40s (starting transient), the system shifts to become *dual-limited*. The presynaptic vesicle number oscillating between zero and near 100 indicates that the system is *vesicle-limited* (top trace). During the downward phase of the vesicle number (top trace), there is a *superimposed ATP limitation*, indicated by oscillation of presynaptic ATP around the threshold value for neuronal firing (dashed line, middle trace). The membrane potential is more depolarized, indicating an UP state (bottom trace). During the upward phase of the vesicle number (top trace), there is an oscillation of ATP (middle trace), but neither ATP nor number of vesicles is limiting. The membrane potential is more hyperpolarized, indicating a DOWN state (bottom trace). Figure 3b shows that after approximately 40s (starting transient), the input frequency alternates between the higher and lower value in sync with the number of vesicles. During the periods of ATP limitation and vesicle limitation (active periods), the presynaptic terminal moving average output frequency *cannot* always match the input frequency; it simply does not have enough resources, either ATP or vesicles or both, to be able to keep up with the input frequency. This is seen as an oscillation around a lower output frequency (top trace) after a brief initial period matching the input frequency (bottom trace). When this occurs, the presynaptic terminal is fatigued and operates only at a level that it can sustain. Because of the influence of the network properties, the system periodically switches into a low-activity rest mode (DOWN state), during which time both input frequency and moving average output frequency take low values (the spontaneous frequency ε), and the presynaptic terminal is able to recover some of its resources. After this recovery occurs, the presynaptic terminal is able to shift back into a high-activity mode for some time before switching back.

**Figure 3.**
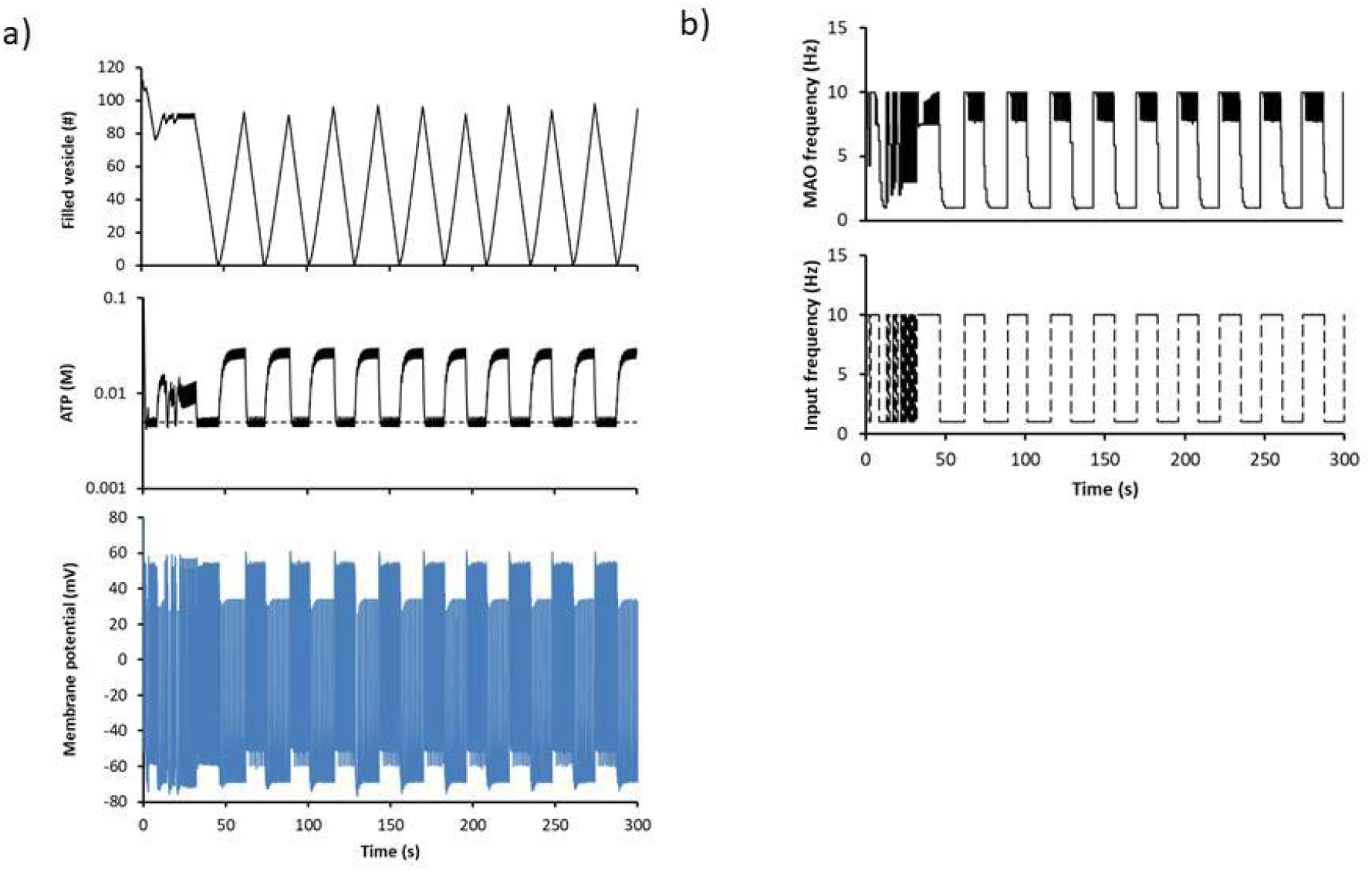
Dual-limited operation. Regime D. *ε* = 1 Hz, *ν* = 9 Hz, *ζ* = 0.7, *χ* = 25 ms. Values shown are for the presynaptic terminal. a) *Top trace*: filled vesicles as a function of time. *Middle trace*: ATP concentration as a function of time. *Bottom trace*: membrane potential as a function of time. b) *Top trace*: moving average output frequency (“MAO frequency”) as a function of time. *Bottom trace*: input frequency as a function of time.

## Limit Cycle Behavior

Simulations at differing input frequencies revealed the presence of limit cycles. Limit cycles represent oscillatory modes of operation in the system. Figure 4 depicts the limit cycles present in four frequency ranges. The starting transient, lasting approximately 40s, has been removed. For clarity in panels (b), (c), and (d), only a single turn of the limit cycle is shown. In Figure 4a, at low input frequencies, the left side of the graph first depicts the running down of vesicles during the first part of the simulation. The limit cycle is described as a tight loop which circulates in a small range with vesicles between one and three (*vesicle-limited* mode; see inset). ATP remains within a tight window of concentrations that lies above the threshold for synaptic vesicle firing (dashed line indicates ATP threshold for firing, 0.005 M). There are two different types of limit cycles in the middle frequency range, which look similar. In Figure 4b, a *vesicle-limited* mode is shown, wherein the number of vesicles drops to zero prior to recovering again—ATP always remains above the threshold for firing (dashed line). Figure 4c depicts *dual-limited* behavior of the system, where there is an *ATP-limited* cycle superimposed on a *vesicle-limited* cycle. This is seen on the left side of the limit cycle, where ATP oscillates to the left and right of the line demarking the ATP firing threshold (below and above the threshold, respectively). Finally, Figure 4d shows an *ATP-limited* regime which occurs at very high frequencies, where the ATP concentration oscillates around the ATP firing threshold, but the number of vesicles never reaches zero.

**Figure 4.**
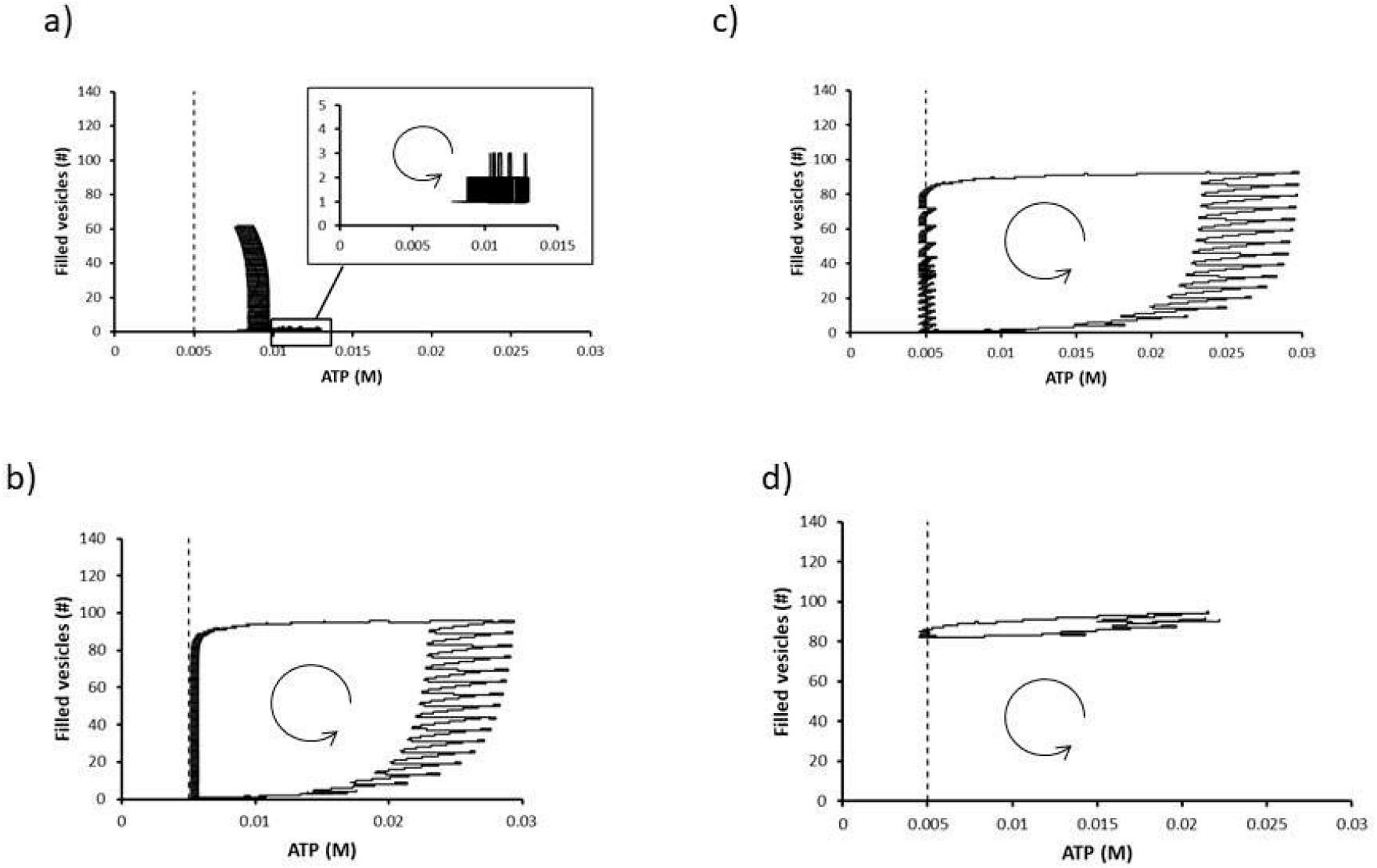
Limit cycle modes. In all figures, the curved arrow indicates the direction of movement as a function of time. In all runs, *ε* = 1 Hz, *χ* = 25 ms. Values shown are for the presynaptic terminal. a) Vesicle-limited mode. Regime B, *ν* = 3 Hz, *ζ* = 0.5 (see Supplementary Discussion). *Inset*: zoomed-in view of the tight loop of limit cycles. b) Vesicle-limited mode. Regime C, *ν* = 7 Hz, *ζ* = 0.5. c) Dual-limited mode. Regime D, *ν* = 9 Hz, *ζ* = 0.7. d) ATP-limited mode. Regime F, *ν* = 22 Hz, *ζ* = 0.4.

The behavior of the presynaptic terminal-astrocyte assembly at different frequencies suggests that there are different regions of operation. These are mapped as functions of the variable component of the input frequency (*ν*) and *ζ* (the proportion of other synapses in the network which need to have the resources to continue to fire) in Figure 5. It is immediately apparent that there are multiple regimes of operation with transition points in between. The input frequency and *ζ* are the two parameters in the model that represent the influence of the rest of the network on the tripartite synapse.

**Figure 5.**
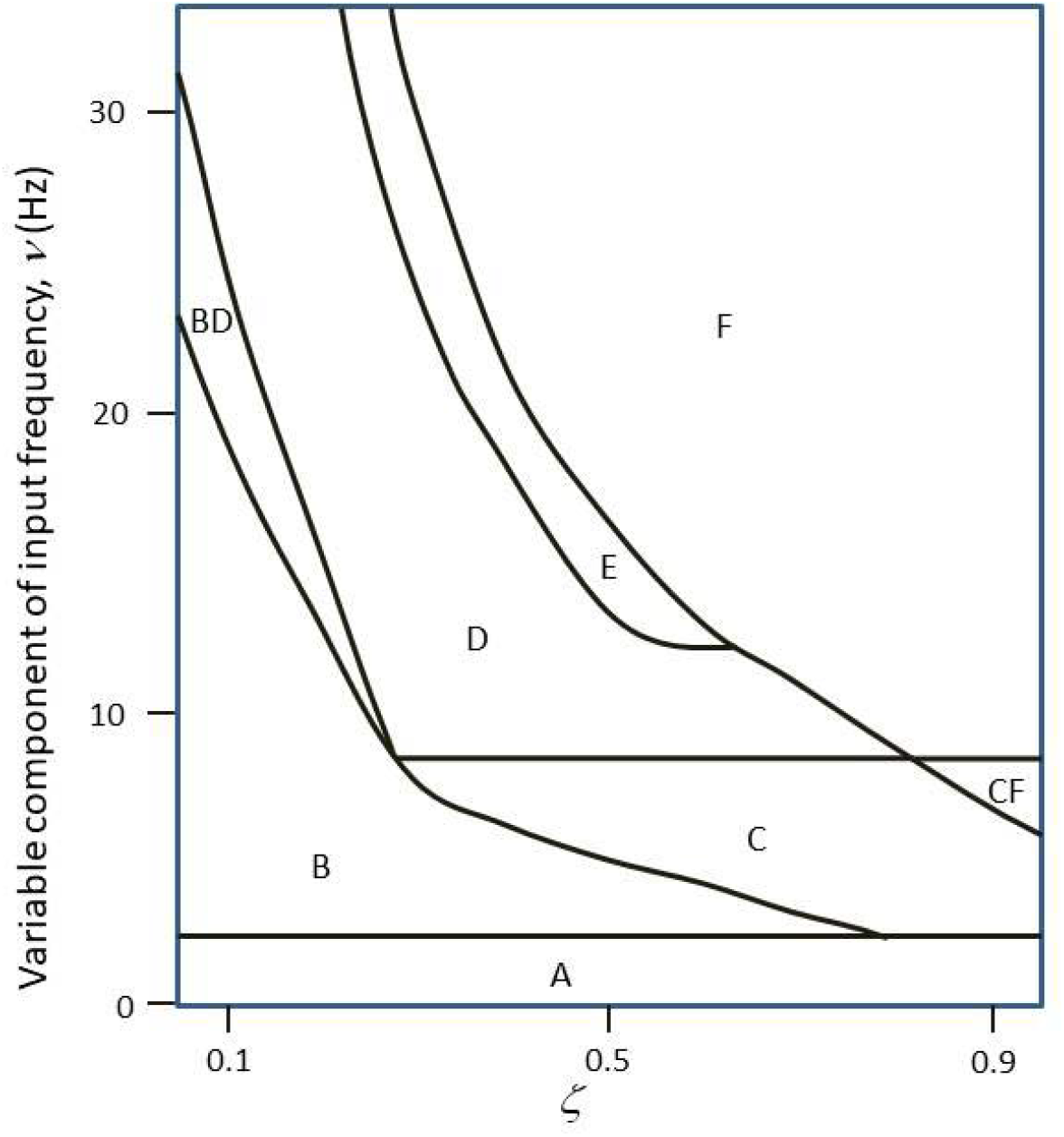
Phase plane analysis. Transitions between regimes were determined by varying the variable component of the input firing frequency (*ν*) and *ζ*. The firing window size (*χ*) was held constant at 25 ms. See Supplementary Discussion for further description of regimes. A: input-limited, B: vesicle-limited (number of vesicles oscillates between 1-3), C: vesicle-limited (number of vesicles has large oscillations), D: dual-limited, E: transitional mode, F: ATP-limited. Regions labeled with two letters are rarer transitional regimes where the behavior of the system initially appears like the second pure regime, but then transitions into the behavior of the first pure regime.

An overview of these regimes is given here; further treatment is found in the Supplementary Discussion. Regime A is the “input-limited” region (Figure 2). There is only one baseline membrane potential (no UP or DOWN states), and the neuron is able to generate an output frequency that is identical to the input frequency. Regime B shows “vesicle-limited” operation (Extended Data Figure 6); there is a small difference in membrane potentials between subsequent firing events, but there are no discrete UP or DOWN states. The number of vesicles oscillates between one and three, and the moving average output frequency oscillates between values that are neither the maximum combined input frequency (*ε* + *ν*) nor the spontaneous frequency (*ε*). Regime C is still a “vesicle-limited” mode (Extended Data Figure 7), however a discrete limit cycle appears, and there are discrete UP and DOWN states indicated by two distinct baseline membrane potentials. Within this region, the number of filled vesicles recovers from zero during the DOWN state. Regime D shows vesicle-limited operation with *superimposed* ATP limitation during the UP state (Extended Data Figures 8 and 9). Both ATP and vesicles recover during the DOWN state in this mode. Regime F is a purely ATP-limited mode (Extended Data Figure 10). Regime E is a transitional mode (Extended Data Figure 11), in which the system starts with purely ATP-limited operation (as in regime F), then transitions to vesicle-limited with superimposed ATP limitation during the UP states (as in regime D).

## Discussion

We have demonstrated a novel model of the tripartite synapse based on chemistry alone, which encompasses the presynaptic terminal and astrocytic process, derived entirely from first principles. Electrical properties, specifically the calculation of the membrane potential, are also based purely on first principles. Our model is robust and flexible, including many important phenomena such as feedback inhibition in regulated metabolic reactions, and diversion of energy substrates into other biosynthetic pathways. Importantly, our model demonstrates “chaotic” features: small changes in a single parameter have far-reaching implications for the behavior of the entire system.

Our model also shows that the presence of rest-activity cycles is a fundamental emergent property of the tripartite synapse. Cycles of high and low activity (“rest”), manifested as limit cycles, intrinsically appear, products of the “chaotic” nature of the system. The model demonstrates use dependence of the output firing frequency—the output firing frequency is dependent on both sufficient ATP and sufficient numbers of filled vesicles, and a high output frequency cannot be sustained without the presence of both. At the higher firing frequencies, the model recapitulates the UP and DOWN states described in the literature^13,18^—though importantly this is due to simulated network effects. The model transitions between the UP and DOWN states depending on the availability of neurotransmitter and ATP, which function as indicators of synaptic fatigue. Through this work, we have shown that energy generation and neurotransmission in the tripartite synapse are intimately linked. This model demonstrates all of the behavioral properties of sleep *at the level of a single tripartite synapse* in a network: 1) quiescence (decreased activity), 2) increased threshold, and 3) homeostasis.^7^ In addition, the model shows that a sleep-like phenomenon is based on and enabled by biochemical processes, and that reduced responsiveness (reduced firing) is necessary for its function (recovery of neuronal resources)—prerequisites for a viable theory of sleep function.^19^

## Supporting information

Supplementary Discussion

## Methods

The model was developed from first principles of chemical reactions in the SimBio environment of MATLAB R2018b (Natick, MA), as a system of 99 ordinary differential equations (see Supplementary Tables). The equations were solved using the ode15s solver in MATLAB. Because neuronal firing is not in a steady-state, using Michaelis-Menten kinetics was inappropriate, and chemical kinetic equations were written using the principle of mass balance, as first order equations. Competitive and non-competitive inhibition for key regulated reactions was modeled as described below.

### Assumptions

#### Key Assumptions

Several key assumptions were made in formulating the model. There were no internal organelles included in either the presynaptic terminal or the astrocyte. This specifically eliminated mitochondria and vesicles as separate compartments, and transport processes between internal compartments and the cytoplasm were eliminated. Because the electron transport chain in mitochondria depends on transfer of electrons inside the mitochondrial membrane, and the generation of ATP ultimately depends on the migration of hydrogen ions across the mitochondrial membrane, this process had to be modeled in an alternative fashion. We elected to model the five complexes of the electron transport chain as follows. Reduced nicotinamide adenine dinucleotide (NADH^+^ + H^+^) reacts with complex 1 to create a reduced complex 1 and an “energy token”. The reduced complex 1 then reacts with complex 2, and further, as well described.^20^ At each point where a hydrogen ion would be transported across the mitochondrial membrane, there is instead the generation of an energy token, such that NADH generates a total of three energy tokens by the end of the electron transport chain. Similarly, reduced flavin adenine dinucleotide (FADH_2_) enters the electron transport chain and generates a total of two energy tokens. The energy tokens then “react” with adenosine diphosphate (ADP) to create ATP at complex 5. Vesicles were also modeled in a simplified fashion. The vesicle loading process was modeled as a single reaction filling a vesicle, and each time the concentration of the “vesicular glutamate” reached a threshold value representing the total amount of glutamate in a full vesicle, the “vesicular glutamate” species was reset to zero concentration, and a counter for vesicles was incremented by one. Finally, for both glycolysis and the TCA cycle, for reactions in which there was no expenditure or generation of energy equivalents (ATP/guanosine triphosphate, NADH, or FADH_2_), such as reactions catalyzed by mutases or isomerases, the product and reactant species were combined into a single species. For example, in the TCA cycle, the conversion of citrate to isocitrate by aconitase does not require any expenditure of energy equivalents, nor does it create any energy equivalents.^20^ In this case, citrate and isocitrate were combined into a single species called “citrate/isocitrate”, and the reaction catalyzed by aconitase was omitted.

#### General Assumptions

Several general assumptions were made regarding the model of the tripartite synapse. Firstly, Michaelis-Menten kinetics was not utilized for any of the enzyme-driven processes, since Michaelis-Menten kinetics assumes a steady-state regime, but the cellular environment in which the processes occur is not in steady-state. Diffusion was assumed to be infinitely fast inside each compartment (there were no intra-compartment diffusion processes modeled). Furthermore, release of glutamate into the synapse was considered instantaneous. Ordinary differential equations were utilized to model diffusion-related losses in the synaptic cleft and ECF. Diffusion was considered to be insignificant between the synaptic cleft compartment and the ECF.

Transport processes in the blood vessel compartment were not modeled, and glucose in this compartment was assumed to have a constant concentration. The post-synaptic terminal was not modeled, and as such, the post-synaptic process of buffering of neurotransmitter via receptor-binding was not considered. Other cellular processes, such as transcription and translation, were not explicitly modeled, but their energy-consuming reactions were lumped together into a “vegetative” process in both the presynaptic terminal and the astrocyte. Finally, the physical processes of synaptic vesicle release, fusion, and reclamation were not modeled; the entire process was assumed to consume a set amount of ATP.

### Assumptions for General Calculations

The values for volumes of the presynaptic terminal, astrocytic process, synapse and vesicle; concentration of neurotransmitter in vesicle; ion fluxes; and intracellular and extracellular ion concentrations were taken from several sources^3,21,22^; some values were derived from others given in these sources. It was assumed that the presynaptic terminal and the astrocytic process both had approximately the same volume as a post-synaptic dendritic spine. Although the ECF compartment is large, in our model, we assumed that the functionally relevant volume of the ECF was approximately the same as the volume of the presynaptic terminal. The range of dendritic spine radius was given as 0.5-1 μm^22^, so we assumed a radius of 0.7 μm. For vesicular size, a range of 40-60 nm in diameter was given^21^; we assumed a diameter of 50 nm. For ion concentrations, there was agreement in most values between Purves *et al^21^* and Attwell and Laughlin^3^, however there was a discrepancy in the intracellular sodium concentrations. Purves *et al^21^* reported a range of 5-15 mM, while Attwell and Laughlin^3^ utilized a value of 20 mM. We thus elected to use a value of 15 mM in our model.

### Assumptions Pertaining to Energy Substrate Metabolism

The following assumptions were made regarding the reactions for energy substrate metabolism. In both the presynaptic terminal and astrocyte compartments, several major side processes—such as fatty acid synthesis and oxidation, and the pentose-phosphate pathway, were not explicitly modeled. Because there was no mitochondrial compartment modeled, the reconversion of citrate into acetyl CoA was not modeled, and fatty acid synthesis was assumed to proceed directly from acetyl CoA. For the presynaptic terminal, the astrocyte-neuronal lactate shuttle (ANLS) hypothesis was assumed^15^, and lactate was the only source of carbon substrate for ATP generation other than conversion from glutamate into α-ketoglutarate. Given some controversy regarding the ANLS hypothesis in the literature (for a review, see Yellen^23^), we did introduce glycolysis into the presynaptic terminal *(data not shown)*, and the five described regions of behavior were recapitulated, with somewhat shifted boundaries due to an increased availability of ATP. However, the model took significantly longer to compute, and the number of data points generated more than quadrupled for the same duration of in-simulation time. As such, we excluded glycolysis in the presynaptic terminal for subsequent runs. For the astrocyte, for simplicity glycogen stores were assumed to be negligible, and neither glycogen synthesis nor glycogenolysis were modeled. A glycogen store would contribute an additional reservoir of energy substrate, which would be expected to have a similar effect to the inclusion of neuronal glycolysis—the behavior of the model would be similar, but with shifted boundaries between the different regimes. We assumed that for both the presynaptic terminal and the astrocyte, the only source of new carbon skeletons for glutamate and glutamine synthesis was conversion from glucose, through the intermediate of α-ketoglutarate, and that sources for amination (but not ATP) were freely available. As with neuronal glucose and astrocytic glycogen above, the addition of a separate source of glutamine would simply contribute an additional reservoir of energy and contribute to the pool of available neurotransmitter, shifting the boundaries of the described regimes.

### Assumptions Regarding Membrane Potentials and Ions

Several assumptions were made regarding ion movements and calculation of membrane potentials. Ion movements were considered only in the presynaptic terminal and ECF compartments; ion fluxes in the astrocyte compartment and synaptic cleft were not considered. Only sodium and potassium ion movements were considered; membrane potential was calculated based on the charge distribution across the presynaptic cell membrane, and a DC (direct current) shift was employed to account for the absence of other ionic species (such as Ca^2+^, Mg^2+^, Cl^-^, HCO_3_^-^, anionic proteins, others) in the model. The important sodium-potassium pump was modeled using three different processes—a process that extrudes sodium out of the cell, a process that brings potassium into the cell, and a process that consumes ATP, for ease of calculation. Of note, because we excluded Ca^2+^ ions, we also excluded the signaling processes relying on calcium. The kinetics of K_ATP_ channels and other specific potassium and sodium ion channels were not explicitly considered. Ion movements occurred during the “firing signal” event (when the presynaptic terminal is triggered to release a synaptic vesicle), and occurred contemporaneously with the vesicular release. Action potentials starting at the axon hillock of the neuron were not modeled, and as such, the Hodgkin-Huxley electrical formalism was not applied. In addition, as noted above, this model primarily reflects chemical processes.

### Determination of Parameters and Initial Conditions

Parameters and initial conditions for the model were selected to yield close-to-physiologic values of all of the species. These parameters and initial conditions are not necessarily the only set of parameters and initial conditions which will yield physiologic values of species, and there are numerous other combinations of values that could yield realistic results. Notably, in the future, many of the parameters can be empirically determined via kinetic experiments in living cells. Nevertheless, there were additional “safeguards” implemented in the model to assure that parameters and species did not attain non-physical values (such as negative values for concentrations, excessively high values for concentrations or negative values for reaction rate constants). Simulation runs were closely examined for evidence of non-physical results. Please see Supplementary Tables for sample values for Compartments, Parameters, Initial Conditions, Events, and Rules. These values represent the parameters for a single run of the model.

### Modeling of Inhibition

Both competitive and non-competitive inhibition were modeled at the appropriate stages in glycolysis and the TCA cycle in the presynaptic terminal and astrocyte. Specifically, for competitive inhibition, the relevant enzyme was considered to be both a product and a reactant in the main reaction. The competitive inhibitor was modeled to bind to the enzyme in a reversible fashion to create an enzyme-inhibitor complex, thereby decreasing the availability of enzyme for the main reaction. For non-competitive inhibition, the reaction rate constants were calculated to be linear functions of the non-competitive (allosteric) inhibitor of the form:

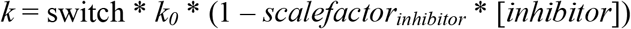

Where *k* is the reaction rate constant, *k_0_* is the ***maximum*** possible rate constant, *scalefactor_inhibitor_* is a scaling factor that determines the strength of the non-competitive inhibition, and *[inhibitor]* is the concentration of the non-competitive inhibitor. *Switch* is a binary variable defined as follows:

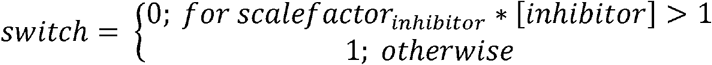

To prevent the reaction rate constant from being negative.

Allosteric activators were modeled similarly, with the special case of cooperativity being modeled by the “activator” being the same species as the substrate:

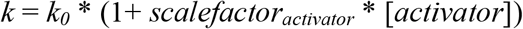

Where *k* is the reaction rate constant, *k_0_* is the minimum possible rate constant, *scalefactor_activator_* is a scaling factor that determines the strength of the allosteric activation, and *[activator]* is the concentration of the allosteric activator.

### Sleep and Ignition

The model has three conditions which determine whether the system is in a high-activity state or a low-activity (rest) state. If the ratio *φ* of output frequency to the combined maximum input frequency (*ε* + *ν*) is greater than *ζ*, the neuron remains in the high-activity state. This first condition indicates that if the presynaptic terminal is able to generate a sufficiently high output frequency, this is a surrogate for the state that a sufficient fraction of the *network* (the other neurons contributing to the input of the presynaptic terminal) also have enough resources to continue in a high-activity period. If *φ* is smaller than *ζ*, which is caused by the output frequency dropping due to limitation of resources (ATP and/or filled vesicles), the neuron drops into a low-activity (rest) state. This second condition indicates that when the presynaptic terminal is unable to generate a sufficiently high output frequency, this is a surrogate for passing a threshold where the *rest of the network* no longer has enough resources to drive a high input frequency to the presynaptic terminal. Once in such a low-activity (rest) state, the neuron would otherwise remain in this state forever, and a third condition functions to switch the system from a low-activity state back into a high-activity state. This occurs when the presynaptic terminal has enough resources to generate a high output frequency. However, because the model is numerically solved with discrete time steps, the output frequency is calculated as a moving average over three time steps (rather than as an instantaneous frequency). In order for the model to “ignite” from a low-activity (rest) state back into a high-activity state, the third condition had to be modified to avoid meeting the conditions for dropping back into the low-activity (rest) state during the first two time-steps before the calculated output frequency has increased to its true higher value. This was accomplished by adding code to the model which held the output frequency equal to the input frequency for the first two steps of “ignition” (when the other two conditions for “ignition”, sufficient ATP and glutamate, were present). The moving average for the output frequency thus effectively increased instantaneously and avoided causing the system to revert back into a low-activity rest state spuriously.

## Data Availability Statement

Data sharing is not applicable to this article as no datasets were generated or analyzed during the current study.

## Code Availability Statement

The Simbio model used for this work is available from the corresponding author upon reasonable request.

## Acknowledgements

This work is not funded by any funding agency, organization, or outside individual. It is not associated with any other work performed at the affiliated institutions.

## Author Contributions

SNJ conceived the work, performed literature review, contributed to the work design, contributed to the analysis of the data, contributed to the manuscript draft, and contributed to the manuscript revisions.

ANJ contributed to the literature review, designed the model, contributed to the analysis of data, drafted the manuscript, and performed manuscript revisions.

NDJ contributed to the work design, analyzed the data, and contributed to the manuscript revisions.

*All authors contributed equally to the work.

## Additional Information

**Supplementary Information** is available for this paper.

## Extended Data

**Extended Data Figure 6.**
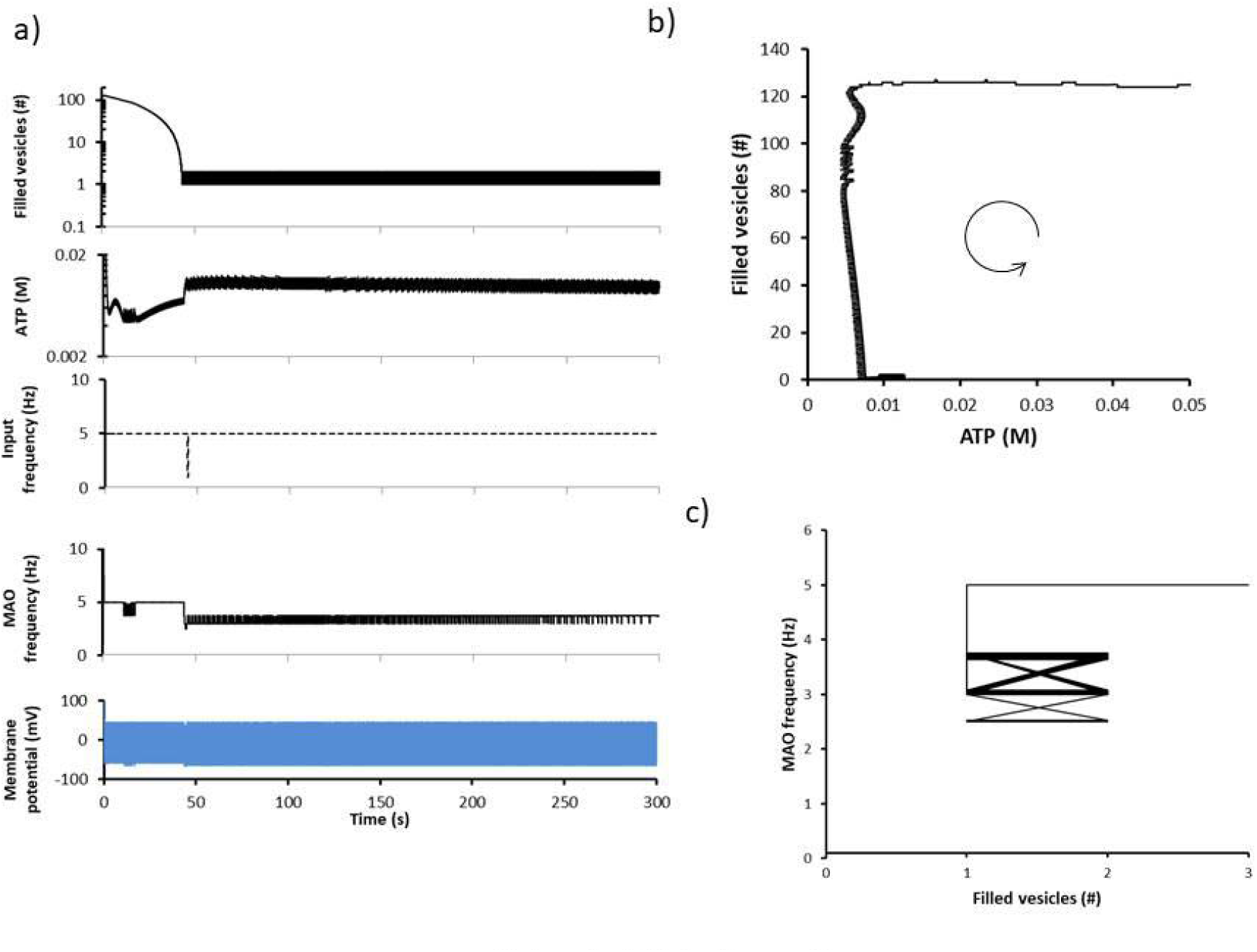
Regime B. *ε* = 1 Hz, *ν* = 4 Hz, *ζ* = 0.5, *χ* = 25 ms. Values shown are for the presynaptic terminal. a) *Top trace*: number of filled vesicles versus time. *Second trace*: ATP versus time. *Third trace*: input frequency versus time. *Fourth trace*: moving average output frequency (“MAO Frequency”) versus time. *Fifth trace*: membrane potential versus time. b) Limit cycle (number of filled vesicles versus ATP). The curved arrow indicates the direction of movement as a function of time. c) MAO frequency versus number of filled vesicles.

**Extended Data Figure 7.**
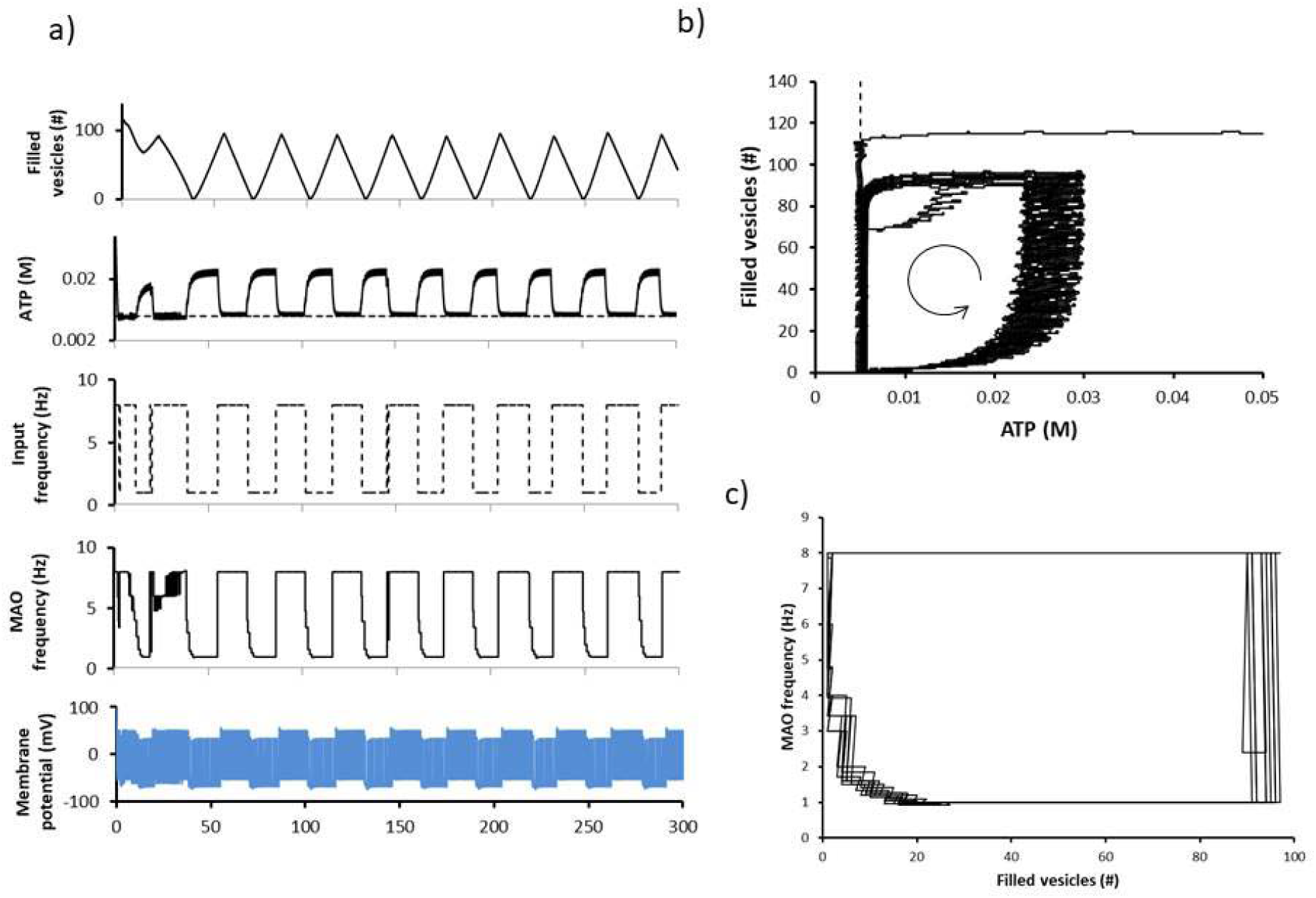
Regime C. *ε* = 1 Hz, *ν* = 7 Hz, *ζ* = 0.5, *χ* = 25 ms. Values shown are for the presynaptic terminal. a) *Top trace*: number of filled vesicles versus time. *Second trace*: ATP versus time. *Third trace*: Input frequency versus time. *Fourth trace*: moving average output frequency (“MAO Frequency”) versus time. *Fifth trace*: membrane potential versus time. b) Limit cycle (number of filled vesicles versus ATP). The curved arrow indicates the direction of movement as a function of time. c) MAO frequency versus number of filled vesicles.

**Extended Data Figure 8.**
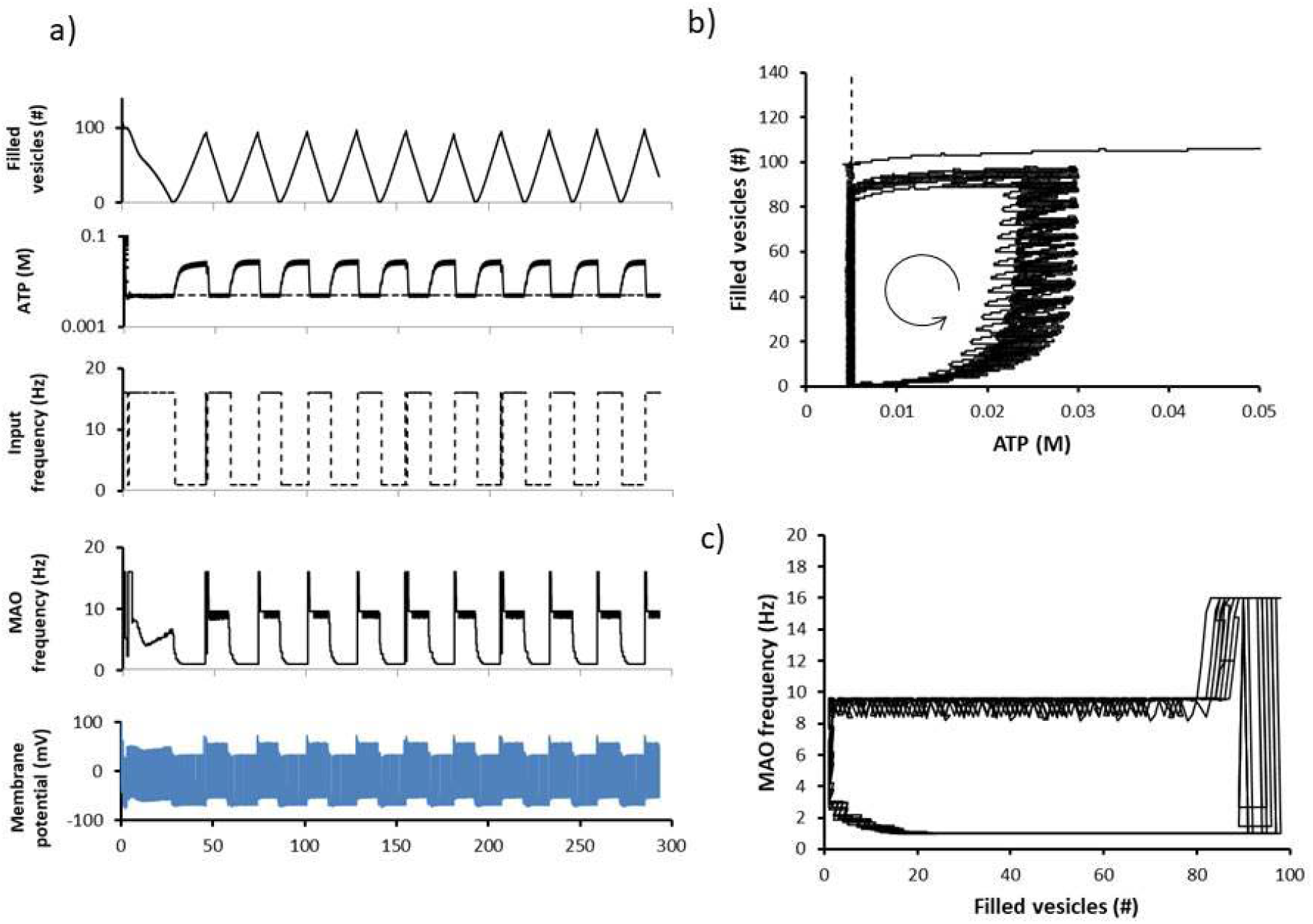
Regime D. *ε* = 1 Hz, *ν* = 15 Hz, *ζ* = 0.2, *χ* = 25 ms. Values shown are for the presynaptic terminal. a) *Top trace*: number of filled vesicles versus time. *Second trace*: ATP versus time. *Third trace*: Input frequency versus time. *Fourth trace*: moving average output frequency (“MAO Frequency”) versus time. *Fifth trace*: membrane potential versus time. b) Limit cycle (number of filled vesicles versus ATP). The curved arrow indicates the direction of movement as a function of time. c) MAO frequency versus number of filled vesicles.

**Extended Data Figure 9.**
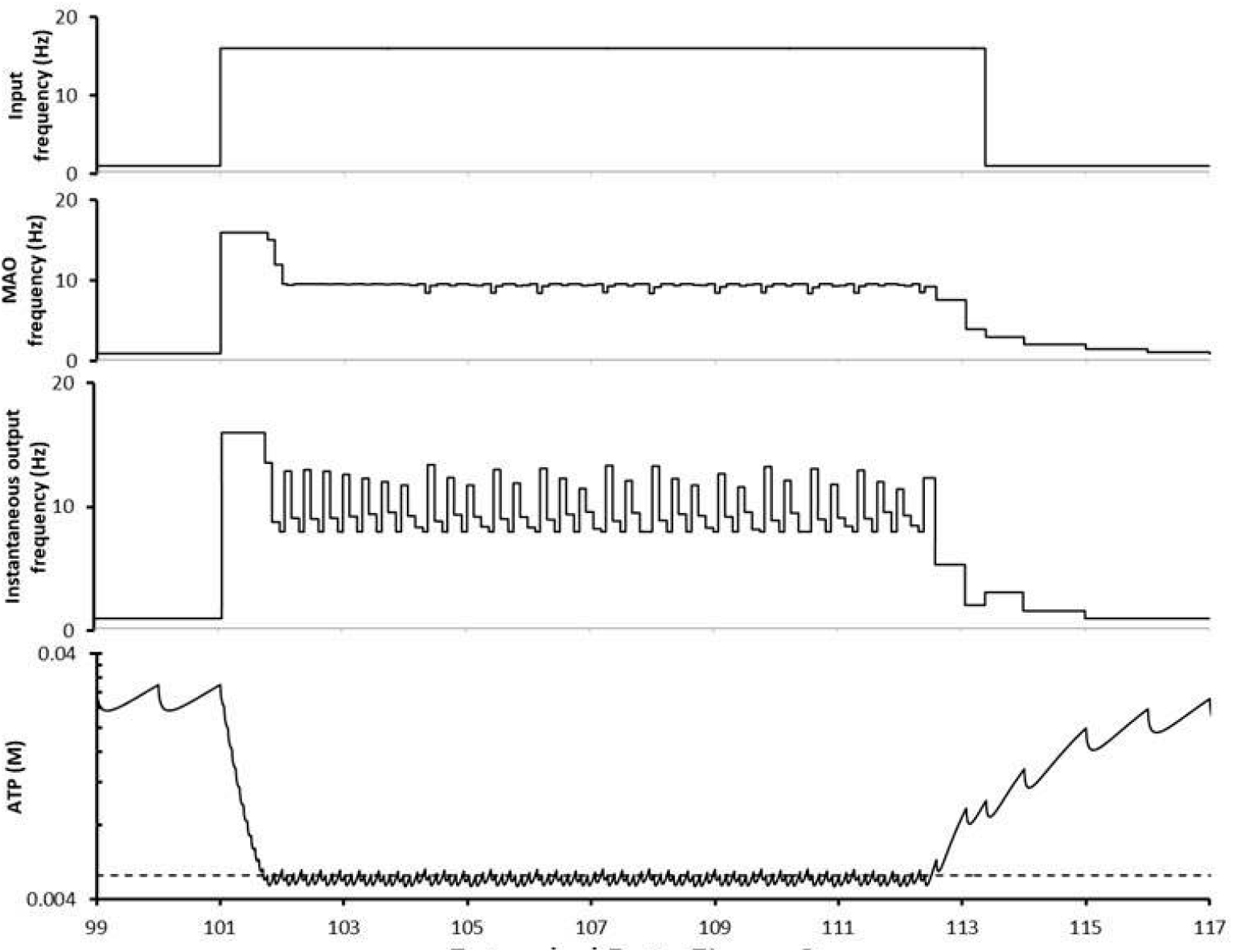
Regime D, a closer view. *ε* = 1 Hz, *ν* = 15 Hz, *ζ* = 0.2, *χ* = 25 ms. Values shown are for the presynaptic terminal. *Top trace*: input frequency versus time. *Second trace*: moving average output frequency (“MAO Frequency”) versus time. *Third trace*: instantaneous output frequency versus time. *Fourth trace*: ATP concentration versus time.

**Extended Data Figure 10.**
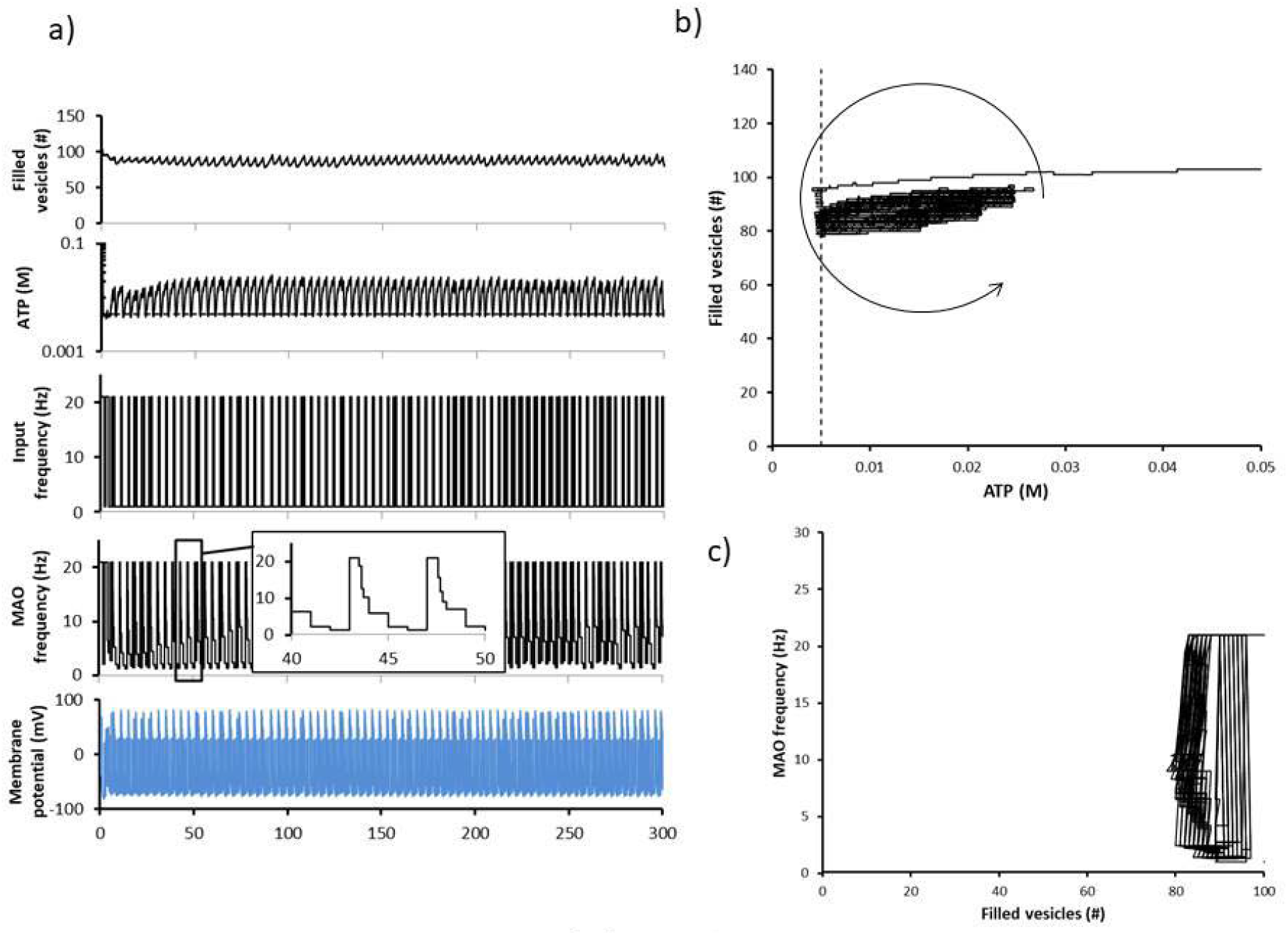
Regime F. *ε* = 1 Hz, *ν* = 20 Hz, *ζ* = 0.5, *χ* = 25 ms. Values shown are for the presynaptic terminal. a) *Top trace*: number of filled vesicles versus time. *Second trace*: ATP versus time. *Third trace*: Input frequency versus time. *Fourth trace*: moving average output frequency (“MAO Frequency”) versus time. *Fifth trace*: membrane potential versus time. b) Limit cycle (number of filled vesicles versus ATP). The curved arrow indicates the direction of movement as a function of time. c) MAO frequency versus number of filled vesicles.

**Extended Data Figure 11.**
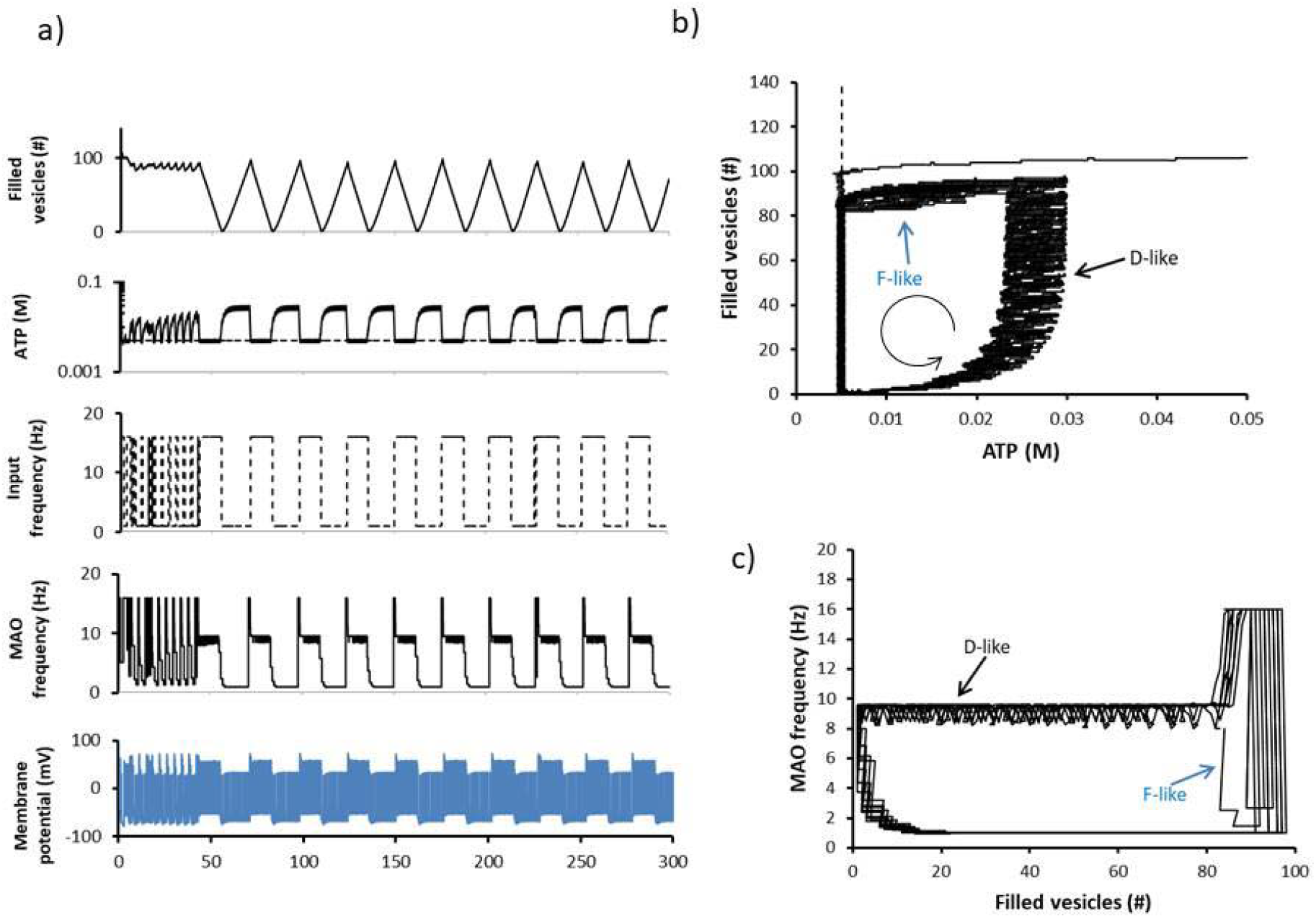
Regime E. *ε* = 1 Hz, *ν* = 15 Hz, *ζ* = 0.5, *χ* = 25 ms. Values shown are for the presynaptic terminal. a) *Top trace*: number of filled vesicles versus time. *Second trace*: ATP versus time. *Third trace*: Input frequency versus time. *Fourth trace*: moving average output frequency (“MAO Frequency”) versus time. *Fifth trace*: membrane potential versus time. b) Limit cycle (number of filled vesicles versus ATP). The curved arrow indicates the direction of movement as a function of time. c) MAO frequency versus number of filled vesicles.

**Extended Data Figure 12.**
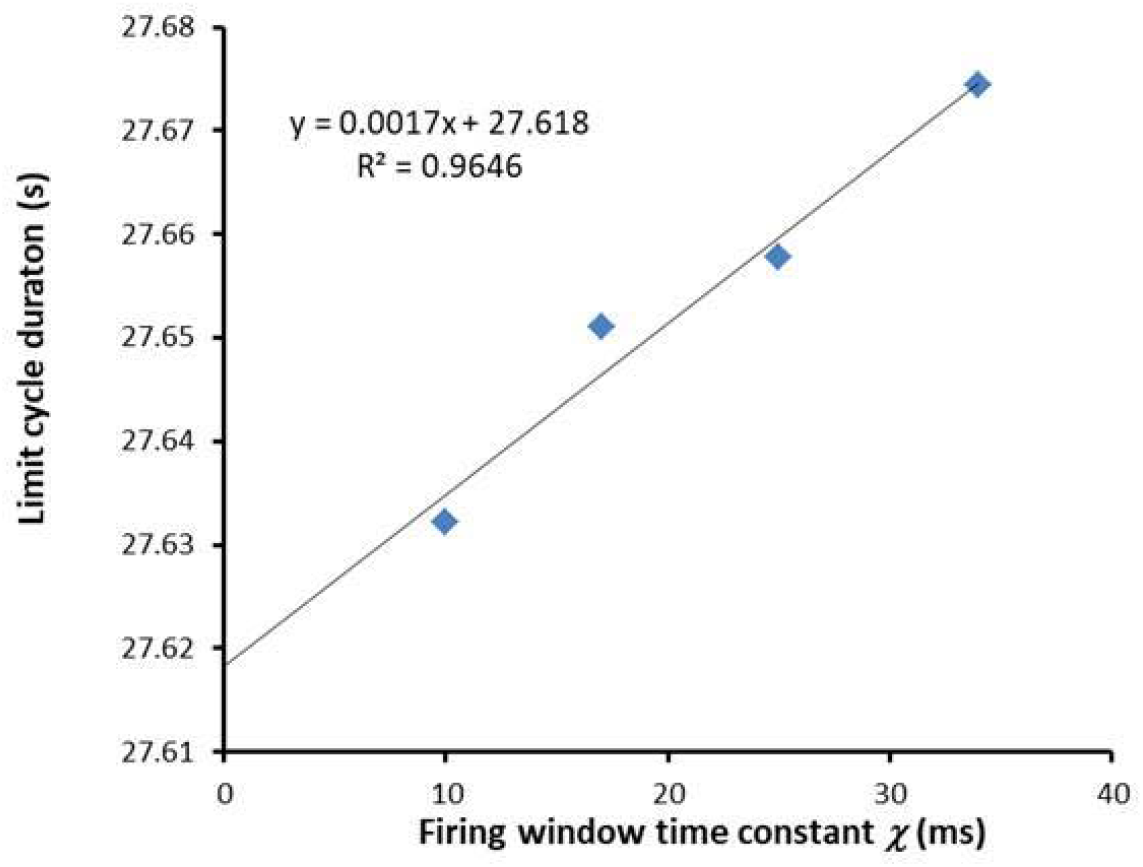
Effect of window size *χ*. The duration of a limit cycle is directly proportional to the time constant *χ*. The y-intercept of the fit line gives what the duration of the limit cycle would be without the confounding effect of using a numerical solver with a maximum allowed step size. *ε* = 1 Hz, *ν* = 8 Hz, and *ζ* = 0.5.

**Extended Data Figure 13.**
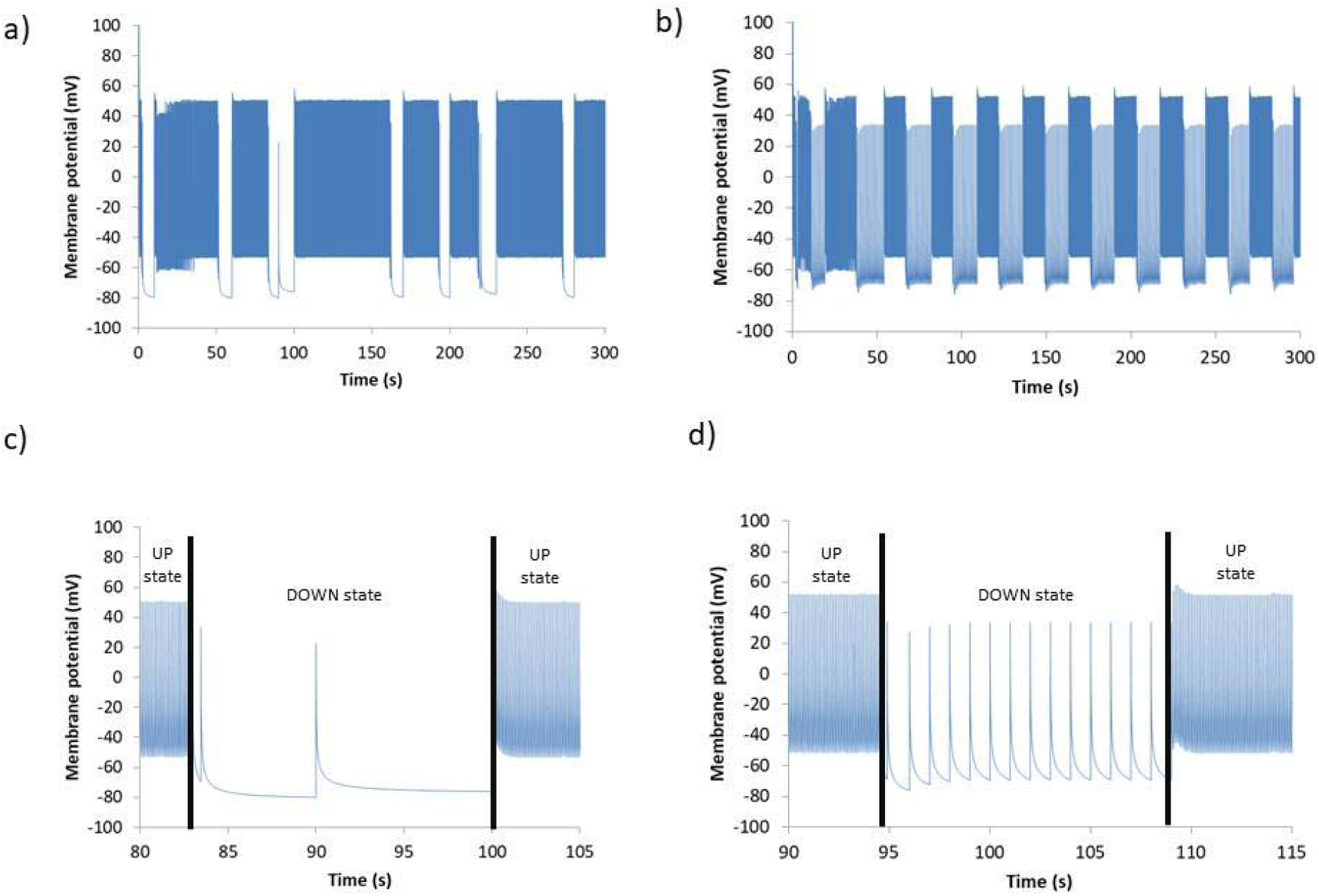
Membrane potentials with varying *ε*. Values shown are for the presynaptic terminal. a) and c) Regime C with *ε* = 0.1 Hz, *ν* = 8 Hz, *ζ* = 0.5, *χ* = 25 ms. The DOWN states are shorter in duration, and the UP states are longer in duration compared to b) and d) Regime C with *ε* = 1 Hz, *ν* = 8 Hz, *ζ* = 0.5, *χ* = 25 ms. In addition, the DOWN states show fewer action potentials at low ε in c), as compared to a higher spontaneous frequency *ε* in d).

**Extended Data Figure 14.**
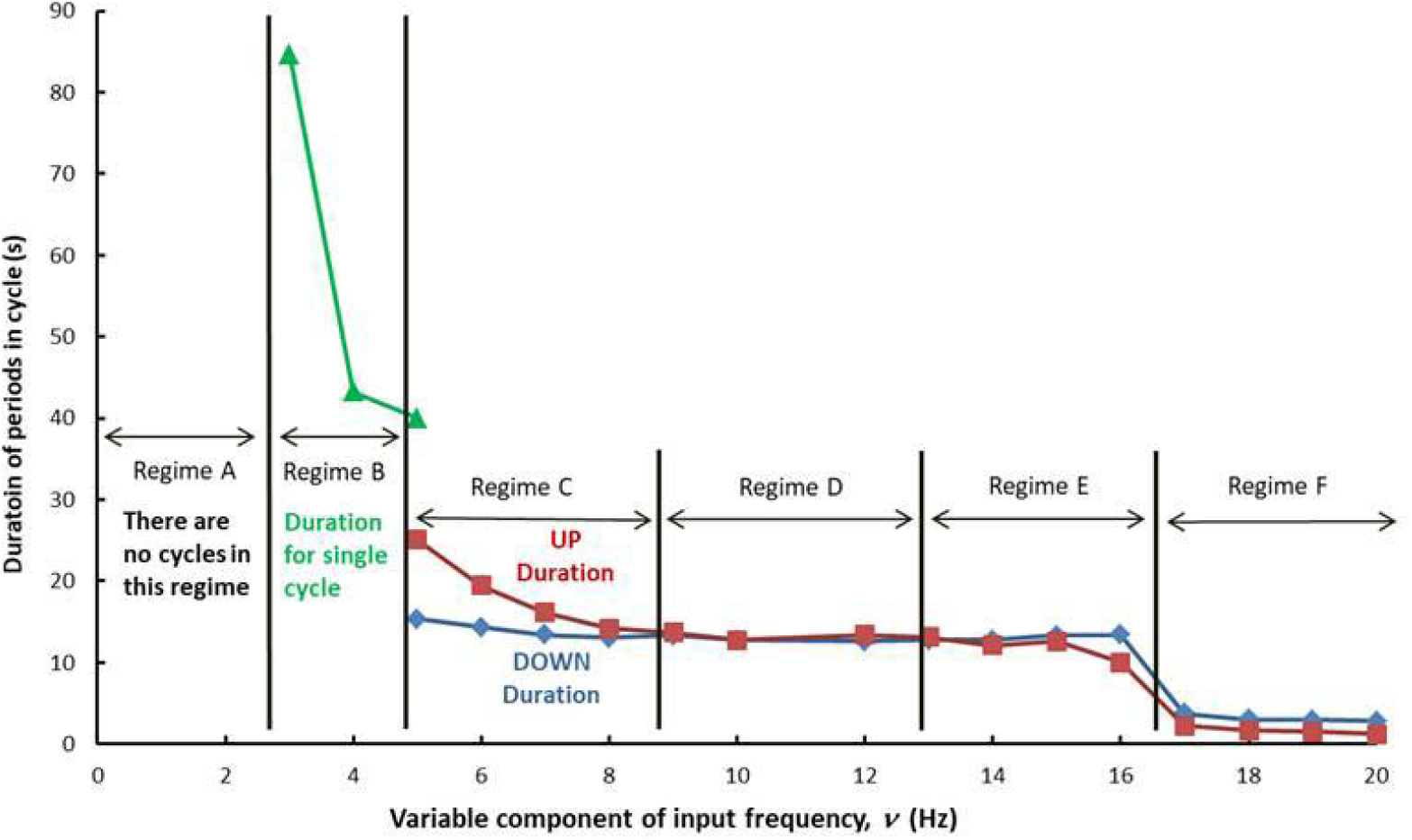
Duration of periods in cycle vs variable component *ν* of input frequency. During regime B, only the duration of the first high input frequency period is plotted. During regime C, the duration of the UP state varies inversely with increasing frequency; it stays relatively stable during regimes D and E. The duration of the DOWN state remains remarkably stable from regime C to regime E. Durations of both the UP and DOWN states drop significantly in regime F. *ε* = 1 Hz, *ζ* = 0.5, *χ* = 25 ms.

## References

1 Liu, C. Y., Yang, Y., Ju, W. N., Wang, X. & Zhang, H. L. Emerging Roles of Astrocytes in Neuro-Vascular Unit and the Tripartite Synapse With Emphasis on Reactive Gliosis in the Context of Alzheimer’s Disease. Frontiers in cellular neuroscience 12, 193, doi:10.3389/fncel.2018.00193 (2018).

2 Farhy-Tselnicker, I. & Allen, N. J. Astrocytes, neurons, synapses: a tripartite view on cortical circuit development. Neural development 13, 7, doi:10.1186/s13064-018-0104-y (2018).

3 Attwell, D. & Laughlin, S. B. An energy budget for signaling in the grey matter of the brain. Journal of cerebral blood flow and metabolism: official journal of the International Society of Cerebral Blood Flow and Metabolism 21, 1133–1145, doi:10.1097/00004647-200110000-00001 (2001).

4 Yu, Y., Herman, P., Rothman, D. L., Agarwal, D. & Hyder, F. Evaluating the gray and white matter energy budgets of human brain function. Journal of cerebral blood flow and metabolism: official journal of the International Society of Cerebral Blood Flow and Metabolism 38, 1339–1353, doi:10.1177/0271678X17708691 (2018).

5 Patel, A. B. et al. The contribution of GABA to glutamate/glutamine cycling and energy metabolism in the rat cortex in vivo. Proceedings of the National Academy of Sciences of the United States of America 102, 5588–5593, doi:10.1073/pnas.0501703102 (2005).

6 McKenna, M. C. The glutamate-glutamine cycle is not stoichiometric: fates of glutamate in brain. Journal of neuroscience research 85, 3347–3358, doi:10.1002/jnr.21444 (2007).

7 Allada, R. & Siegel, J. M. Unearthing the phylogenetic roots of sleep. Current biology: CB 18, R670–R679, doi:10.1016/j.cub.2008.06.033 (2008).

8 Verkhratsky, A. & Nedergaard, M. Physiology of Astroglia. Physiological reviews 98, 239–389, doi:10.1152/physrev.00042.2016 (2018).

9 Jolivet, R., Coggan, J. S., Allaman, I. & Magistretti, P. J. Multi-timescale modeling of activity-dependent metabolic coupling in the neuron-glia-vasculature ensemble. PLoS computational biology 11, e1004036, doi:10.1371/journal.pcbi.1004036 (2015).

10 Shen, J. et al. Determination of the rate of the glutamate/glutamine cycle in the human brain by in vivo 13C NMR. Proceedings of the National Academy of Sciences of the United States of America 96, 8235–8240, doi:10.1073/pnas.96.14.8235 (1999).

11 Magistretti, P. J. Neuron-glia metabolic coupling and plasticity. The Journal of experimental biology 209, 2304–2311, doi:10.1242/jeb.02208 (2006).

12 Belanger, M., Allaman, I. & Magistretti, P. J. Brain energy metabolism: focus on astrocyte-neuron metabolic cooperation. Cell metabolism 14, 724–738, doi:10.1016/j.cmet.2011.08.016 (2011).

13 Jercog, D. et al. UP-DOWN cortical dynamics reflect state transitions in a bistable network. eLife 6, doi:10.7554/eLife.22425 (2017).

14 Engl, E. & Attwell, D. Non-signalling energy use in the brain. The Journal of physiology 593, 3417–3429, doi:10.1113/jphysiol.2014.282517 (2015).

15 Pellerin, L. & Magistretti, P. J. Sweet sixteen for ANLS. Journal of cerebral blood flow and metabolism: official journal of the International Society of Cerebral Blood Flow and Metabolism 32, 1152–1166, doi:10.1038/jcbfm.2011.149 (2012).

16 Schousboe, A., Waagepetersen, H. S. & Sonnewald, U. Astrocytic pyruvate carboxylation: Status after 35 years. Journal of neuroscience research 97, 890–896, doi:10.1002/jnr.24402 (2019).

17 Ramirez, D. M. & Kavalali, E. T. Differential regulation of spontaneous and evoked neurotransmitter release at central synapses. Current opinion in neurobiology 21, 275–282, doi:10.1016/j.conb.2011.01.007 (2011).

18 Vyazovskiy, V. V. & Harris, K. D. Sleep and the single neuron: the role of global slow oscillations in individual cell rest. Nature reviews. Neuroscience 14, 443–451, doi:10.1038/nrn3494 (2013).

19 Krueger, J. M., Frank, M. G., Wisor, J. P. & Roy, S. Sleep function: Toward elucidating an enigma. Sleep medicine reviews 28, 46–54, doi:10.1016/j.smrv.2015.08.005 (2016).

## Methods References

20 Berg, J. M., Tymoczko J.L., and Stryer L. Biochemistry. 6th edn, (WH Freeman and Company, New York, 2007).

21 Neuroscience. 4th edn, (Sinauer Associates, Inc., Sunderland, MA, 2008).

22 Fiala J.C., H. K. M. in Dendrites (ed Spruston N. Stuart G., and Hausser M.) 1–34 (Oxford University Press, Inc., New York, 1999).

23 Yellen, G. Fueling thought: Management of glycolysis and oxidative phosphorylation in neuronal metabolism. The Journal of cell biology 217, 2235–2246, doi:10.1083/jcb.201803152 (2018).

