## Supplementary Discussion for "Sleep-like behavior is a fundamental property of the tripartite synapse"

#### Motivation for Sleep

##### What is sleep and why is it necessary?

The nature and function of sleep have eluded philosophers and scientists for centuries, and even a single, cohesive definition of what constitutes sleep is difficult, although many definitions and functions have been postulated over time. We will here summarize some of the main descriptions and postulated purposes for sleep.

##### Organism Level

At the level of the organism, several behavioral characteristics are felt to indicate that an organism is "sleeping", or in a "sleep-like" state. Diverse species have been observed to assume a particular posture and close their eyes. They also have an increased threshold for activity, meaning that they require a stronger stimulation to react to a sensory stimulus than they would during a wakeful state.[24] This low-activity state is accompanied by changes in various physiological factors, including muscle activity, heart rate, and brain temperature.[25] In addition, during certain parts of sleep, there is a loss of thermoregulation in endotherms.[26]

In addition, variants of sleep have been observed which challenge both the typical definitions of sleep at a behavioral level and an electrical level (see below). Some species, such as cetaceans and fur seals, have been observed to have unihemispheric sleep—behaviorally they have decreased activity on one side of the body, which correlates with electroencephalographic evidence of sleep in the contralateral hemisphere.[27] Other species, particularly migratory birds, have "micro-sleeps", wherein they behaviorally sleep for seconds at a time *while flying*.[28] There is also evidence that the brain can even have local sleep: when rats' whiskers were repetitively stimulated, the rats became progressively less responsive to further stimulation of the same whisker (increased behavioral threshold), and (only) the corresponding barrel cortex showed the electrographic equivalent of sleep.[29] Species as simple as the *C. elegans* and the jellyfish *Cassiopea* have been shown to have alternating periods of high activity ("wake-like") and low activity ("sleep-like") states[30,31], suggesting that sleep is a universal and necessary phenomenon in sufficiently complex organisms. Proposed functions of sleep at this level include conservation of energy, (relative) safety from predation, and immune function.[19,32] Interestingly, the likely presence of numerous combinations of parameters yielding physiological behavior of neurochemical species evident in our model may explain the vast diversity of sleep behaviors across animal species.

Brain Level

At the level of the brain, in mammals, sleep has been defined by the presence of distinctive changes in the electroencephalogram (EEG). There is an orderly progression of changes in the patterns of EEG, which has been found to be shared in multiple species. Sleep is characterized as having two main components: non-rapid-eye-movement (NREM) sleep and rapid-eye-movement (REM) sleep.[25] A great deal of research has been done to elucidate the networks that are active during the NREM and REM sleep states, which are thought to be responsible for transitioning between the different states; these will be reviewed below. However, significantly less is known about how the activity of individual neurons in the sleeping cortex generates the sleep potentials recorded by scalp EEG. Some groups have shown *in vivo* that behavioral NREM sleep correlates with alternating periods of relatively less hyperpolarized membrane potentials with a burst firing pattern (the UP state) with periods of relatively more hyperpolarized membrane potentials with absence of firing (the DOWN state).[13,18] Others have suggested that thalamocortical oscillations are responsible for some EEG features of sleep, including cortically-generated slow waves and thalamically-generated spindles.[33] We are not aware of any studies that attempt to describe the cellular level firing patterns that correspond to the low amplitude fast activity seen during REM sleep, but this activity is generally felt to be similar to the activity seen during wakefulness.[33] At the level of the brain, there are descriptions of what constitutes sleep, but there do not seem to be clear functions for it that are distinct from those noted at the behavioral level.

Cellular Level

Sleep is challenging to define at the cellular level; however, it has been proposed that the cortical column, or a different neuronal assembly, is the basic functional module of the cortex, and that these columns can "sleep". In this context, "sleep" has been suggested to be defined as decreased activity (decreased firing rate), increased threshold (manifested as more hyperpolarized membrane potentials), and use dependence (fatigability).[34] As noted above, cortical columns in rat barrel cortex have been shown to have these characteristics *in vivo*. Distinctive patterns of local field potential (LFP) activity were also found which correlated with the high-activity wake-like states and the low-activity sleep-like states.[29]

At the cellular level, low-activity sleep-like periods have been proposed to have multiple possible functions including synaptic pruning, synaptic stabilization, protein synthesis, and other synaptic maintenance activities.[18] These cellular-level functions may mediate higher-level functions including learning and memory consolidation.[19]

How is sleep related to metabolism and energy?

Fundamentally, it is commonly accepted that organisms sleep when they are "tired", which is to say, when they have run out of resources. This then implies that one of the functions of sleep is for the brain to switch into a mode of operation that conserves energy, as noted above.[19]

Although there is some variability between studies, several groups have found that brain metabolism is in fact decreased during NREM sleep, compared to wakefulness. On the other hand, it has been found that brain metabolism during REM sleep is high.[26] Some groups have proposed that REM sleep functions as a way for the brain to shift back into a waking state from an NREM sleep state.[35] As such, the REM state necessarily would have higher metabolic demands than NREM sleep, and would approximate the energy demands of the waking state.

How is sleep related to our model?

Neuronal/glial networks grown *in vitro* have been shown to display some properties suggestive of organism-level sleep. This finding highlights the difference between two major classes of theories of sleep—that sleep is to maintain synaptic connectivity (a network-level explanation), and that sleep is to restore energy sources (a cellular-level explanation). It has been pointed out that these two classes of theories are not mutually exclusive and that "sleep could be a cellular property precisely because the cell's biochemistry, including its metabolism, is driven by network activity".[19] We agree with this latter view, and thus took as a hypothesis that the tripartite synapse, driven by network input, could demonstrate the basic features of sleep—decreased activity, increased threshold, use-dependence, and homeostasis.[7] However, it is clear that the tripartite synapse does not exist in isolation; it is part of a network of other, similar tripartite synapses. Groups have shown that neurons in isolation *in vitro* fire in a low frequency "sleep-like" pattern, but they switch into a more complex high frequency "wake-like" state when stimulated externally via neurotransmitter or electricity, and they cycle between "sleep-like" and "wake-like" states.[34] The external inputs simulate the network effects that the neurons participate in *in vivo*. The investigators themselves perform (some of) the functions of the astrocytes, clearing metabolites and delivering nutrients by means of medium changes, thus completing the simulated tripartite synapse. The rudiments of "sleep", then, are seen at the level of the individual tripartite synapse, participating in a network with other tripartite synapses. Collections of interconnected tripartite synapses comprise cortical columns and other networks that generate sleep-like behavior, but interestingly, there is no definite requirement for either a minimum number of neurons or a specific configuration of connectivity.

We further propose that networks of cortical columns (or similar networks) also generate cycles of rest and activity at a different time scale. These cycles-within-cycles of rest and activity at different levels of organization generate the phenomenon called "sleep" at the level of the organism. The oscillatory dynamical behavior of the tripartite synapse determines the emergence of sleep at higher levels of organization (cortical columns, networks of columns, functional networks at the regional-lobar level, and at the brain level), but sleep cannot and should not be reduced to simply a summation of the behavior of all of the tripartite synapses in the brain.

### Comparison to other models

We will begin with a brief survey of neurochemical models of neurons, with or without the addition of glia, vasculature, or other elements. We will then present a few of the published electrical models of neurons, which are relevant to modeling sleep. Finally, we will explain the rationale for choosing a neurochemical model for this work.

#### Neurochemical Models

There are only a few studies that investigate neurochemical models of the tripartite synapse (defined as systems that model the biochemical reactions that take place in the different compartments of the tripartite synapse). Jolivet *et al* proposed a biophysical model including a neuronal compartment, astrocytic compartment, extracellular compartment, and a vascular compartment. They extended prior work and generated a set of 33 differential equations, with compartmentalization between cytosolic and mitochondrial compartments in the neuron and the astrocyte, the inclusion of a Hodgkin-Huxley-type electrical model of synaptic firing, the provision of explicit glutamatergic input, and explicit calculation of sodium transport. Kinetics related to Michaelis-Menten kinetics was utilized to represent some enzyme-driven processes. Using this model, they were able to reproduce the evoked responses of various parameters seen in rat and human *in vivo* experiments.[9] Patel *et al* examined the relationship between glutamatergic neurotransmission and neuronal glucose oxidation in a seizure model using $^{13}C$ nuclear magnetic resonance (NMR). As part of their study, they established a metabolic model to calculate $^{13}C$ fluxes in the neuron and astrocyte, based partly on Michaelis-Menten kinetics. They found that neuronal activity, neurotransmitter cycling, and glucose oxidation were linearly coupled over much of the range of neuronal activity.[36]

Electrical Models

Many studies have looked at electrical models of neuronal action potential firing; most of these have used the Hodgkin-Huxley formalism or a variant of the same. We will briefly review several of the studies that examine the interplay between neurons and astrocytes using an electrical model. Tewari and Majumdar explicitly modeled a tripartite synapse to evaluate the influence of astrocytes on synaptic plasticity. They found that astrocytic calcium fluxes enhanced synaptic augmentation in their model, demonstrating the importance of including the astrocyte in a model of synaptic transmission.[37] Flanagan *et al* looked at the effect of astrocytic glutamate handling on post-synaptic neuronal excitability. They found that intracellular astrocytic glutamate concentrations substantially affect the behavior of synaptic transmission.[38] Overall, numerous studies have shown the importance of the interaction between neurons and astrocytes using models derived from the Hodgkin-Huxley formalism. One study[39] attempted to more directly connect metabolic considerations to the Hodgkin-Huxley formalism by introducing an

ATP-gated current mediated by the $K_{ATP}$ channel. They found that the model was able to produce a pattern consistent with "burst suppression", wherein neurons show bursts of activity with intervening periods of quiescence. We are not aware of any other models other than that in Jolivet *et al*[9] which try to connect the neurotransmission of neurons with their metabolic processes.

Why a Neurochemical Model?

We chose to investigate the connection between neurotransmission and metabolism from a purely neurochemical approach. While the electrical models, exemplified by the Hodgkin-Huxley formalism, have successfully been used to model various neuronal and astrocytic signaling processes, the electrical phenomena that are modeled are driven by underlying neurochemical processes. These underlying processes are completely abstracted away when using an electrical model, thereby breaking any direct linkages with other metabolic processes such as energy generation and usage, protein synthesis, or cellular housekeeping functions. In their seminal paper advancing the original mathematical form of the eponymous formalism, Hodgkin and Huxley expressly stated that the agreement of their equations with experimental data should not be taken to mean that the equations were anything more than an empirical description of the observed phenomenon. They further noted that although there were possible physical interpretations for certain features of their equations, the predictive success of the equations did not mean that the associated physical interpretation was correct.[40] The advantage of a neurochemical model is that it more directly models the underlying physiological processes, and is also flexible enough to account for other effects such as competing reactions, spatial considerations, and even thermodynamic effects. A neurochemical model can comprehensively represent the metabolic processes and the electrical properties can be derived out of this. The rapid increase in computational power over time has made it possible to model a large number of metabolic processes numerically, something which was not practically doable even five years ago. Our work represents the first of a potential world of robust, detailed, increasingly physiologically accurate neurochemical models that can help explain the workings of brain processes.

Window Effects

We also investigated the effects of varying the window size $\chi$ on our model. The model was run varying $\chi$ from 10-34 ms (corresponding to maximum neuronal firing frequencies of 100 to 30 Hz), and duration of limit cycle versus window size was plotted (Extended Data Figure 12). Interestingly, the duration of the limit cycle generated was a linear function of $\chi$, with an $R^2$ of 0.9646. The line could be extrapolated to a window size of zero. This gives a value for the duration of the limit cycle removing the confounding effect of the maximum allowed step size for the discrete numerical solver with the window size. Window size is important, because it determines whether or not the presynaptic terminal will actually fire. The firing window also determines the maximum possible frequency that the neuron can fire ($1/\chi$); otherwise firing windows would overlap.

Network Effects on Rest-Activity Cycles

Components of the Input Frequency: $\varepsilon$ and $v$

As noted previously, the input frequency and $\zeta$ are the two parameters that represent the influence of the rest of the network on the presynaptic terminal. The maximum value that the input frequency ever takes has been called the “combined maximum input frequency”, and is equal to the sum of a constant component ($\varepsilon$) and a variable component ($v$). The values of $v$ determine which regime the system operates in, as described earlier. The values of $\varepsilon$ also change the way the system runs, though in a somewhat different way—specifically, they determine how long DOWN states last. For example, if $\varepsilon$ is smaller (Extended Data Figure 13a for lower $\varepsilon$, compared to Extended Data Figure 13b for higher $\varepsilon$), this means that the presynaptic terminal fires less frequently *spontaneously* during the DOWN state—seen best in the zoomed-in views of small ε (Extended Data Figure 13c) and larger ε (Extended Data Figure 13d). If the frequency of spontaneous firing ($\varepsilon$) is lower, the presynaptic terminal uses less ATP and vesicles for firing events, and will recover its resources faster, leading to a shorter DOWN state (Extended Data Figure 13a compared to Extended Data Figure 13b). The UP state duration becomes longer as well, since the combined maximum input frequency ($\varepsilon + v$) is also smaller, but this effect is less pronounced since usually $v >> \varepsilon$.

Our model also reproduces the temporal changes in membrane potentials seen in individual neurons during slow wave sleep. In intracellular recordings in naturally awake and sleeping animals, it was shown that during a slow wave sleep state, individual neurons had short bursts of spiking with relatively high inter-burst membrane potentials, punctuated by periods of very hyperpolarized resting membrane potentials.[41] The membrane potential during the transition period between these fast bursts (UP states) and hyperpolarized states (DOWN states) decreased exponentially. A similar transition between the UP state and DOWN state occurs in our model without utilizing the Hodgkin-Huxley formalism (Extended Data Figures 13c and Extended Data Figure 13d). Notably, the membrane potential looks particularly similar to the *in vivo* UP and DOWN states for the smaller value of ε, at 0.1 Hz (Extended Data Figure 13c).

$\zeta$ and $\varphi$

The input firing frequency is a function of the ratio of the moving average output frequency of the presynaptic terminal to the combined maximum input frequency ($\varepsilon + v$)—called $\varphi$. In addition, we have defined a threshold value $\zeta$, which represents the fraction of other synapses in the network that must have sufficient resources available in order for the network to continue firing at a high rate. This is an indirect measure of the connectivity between neurons. If the value of $\varphi$ falls below the value of $\zeta$, then the network switches into a low spontaneous firing frequency mode; this functionally means that the *input frequency* drops to a low baseline frequency representing the sum total of *spontaneous* inputs from the rest of the network. The resultant moving average output frequency is then the outcome of spontaneous action potentials generated by the low-level network inputs, and the presynaptic terminal's own spontaneous neurotransmitter release. If the presynaptic terminal later builds sufficient ATP and vesicles, then the network switches back into a high frequency mode determined by an input firing frequency equal to the spontaneous firing frequency ($\varepsilon$) + the independent parameter of the variable component of the input frequency ($v$).

The relationship between $\zeta$ and $\varphi$ will be explained further—while both quantities are characteristic numbers in the system, $\zeta$ is an independent parameter and $\varphi$ is a variable quantity. The condition when $\varphi$ is less than $\zeta$ represents a situation in which the presynaptic terminal is unable to generate a threshold fraction of synaptic release events over time (moving average output frequency). When this threshold is reached, the inequality $\varphi < \zeta$ is taken as a surrogate for the condition that a fraction ***at least*** $(1 - \zeta)$ of *other* presynaptic terminals participating in the network are *also* unable to generate the threshold fraction of synaptic release events over time. This is the reason for the input frequency to drop to the low spontaneous rate. Similarly, when the presynaptic terminal recovers enough resources that it has more than a threshold value of ATP and vesicles, this threshold also is a surrogate for the recovery of resources in the other presynaptic terminals in the network such that ***no more than*** $(1 - \zeta)$ other presynaptic terminals are depleted of resources. The system is then “ignited” such that $\varphi > \zeta$ is again true. As such, $\zeta$ represents a critical number which describes the point at which the entire network switches between a high frequency and a low frequency mode.

The quantities $\zeta$ and $\varphi$ define four possible states for the network. The first state is when both the presynaptic terminal of interest and the network have sufficient ATP and available neurotransmitter, leading to a high input and output firing frequency at the presynaptic terminal. In this case, $\varphi > \zeta$. The second state occurs when both the presynaptic terminal and the network have insufficient ATP or available neurotransmitter to be able to function at a high frequency. Neither can generate high frequency firing, and $\varphi < \zeta$. The third state is produced when the presynaptic terminal of interest has built up a sufficient supply of ATP and available neurotransmitter to be able to fire again at a high rate, but the network is not yet firing at a high rate. In this situation, the input frequency to the presynaptic terminal is initially low, but the network becomes “ignited” because the presynaptic terminal is able to fire at a high output frequency again ($\varphi > \zeta$). These three states correspond to the three conditions described in “Sleep and Ignition” above. There is a fourth possible state, when the presynaptic terminal of interest does not have enough resources to continue to fire at a high rate, but the rest of the network does ($\varphi < \zeta$). This state is not represented in our model, since we hold that when the presynaptic terminal of interest is depleted of resources, this implies that the rest of the network also does not have enough resources (the second state described above). However, this fourth state would replicate a form of “short term depression” caused by the inability of the neuron to release synaptic vesicles[42], due to either a lack of ATP or a lack of filled vesicles.

In addition to functioning as the threshold for switching between states, the quantity $\zeta$ also determines the boundaries between the different modes of firing for the presynaptic terminal.

Looking back at Figure 5, it can be seen that there are curves that separate the different modes of operation of the system. The modes of operation, or regimes, will be described further in a following section. Regimes A, B, C, D, and F may be considered the five primary firing modes of the presynaptic terminal. Regime C is similar to regime D; however in regime C, the output frequency is always equal to the input frequency in the UP state, whereas in regime D, the output frequency is mostly lower than the input frequency. Regime E appears to be a hybrid mode with features of regimes D and F. At the boundaries between regimes, there are also several other hybrid modes that start in one regime and end in another, such as BC (first appears like C and then transitions to B), BD (first appears like D, and then transitions to B), and EF (first appears like F, then transitions to E). The regime A to regime B transition appears to be independent of $\zeta$, however it is possible that there is some minimal dependence that is not captured by the sampling of $\zeta$ values. The regime B to regime C/D transition frequency is described by a curve:

$$\nu_{BCD} = \frac{2.7032}{\zeta^{1.009}}$$

Where $\nu_{BCD}$ is the value of the variable component of the input frequency (an independent parameter, $\nu$) where the transition from regime B to regime C/D occurs. This curve fits extremely well with the transition frequency locations, and has an $R^2$ value of 0.986. Interestingly, the transition frequency is very close to a multiple of the inverse of $\zeta$. There is again a demarcation between regimes C and D which does not appear to be dependent on $\zeta$, although a minimal dependence (not captured by the sampling of $\zeta$ values) cannot be excluded. Notably, at low values of $\zeta$, regime C disappears, and there is a transition from regime B, to hybrid regime BD, directly to regime D. The transition between regimes D and E/F can be described by a curve:

$$\nu_{DEF} = \frac{7.1327}{\zeta^{1.23}}$$

Where $v_{DEF}$ is the value of the variable component of the input frequency (an independent parameter, $v$) where the transition from regime D to regime E/F occurs. This curve also tightly fits the transition frequency locations, and has an $R^2$ value of 0.993. In this case as well, the transition frequency is close to a multiple of the inverse of $\zeta$.

What are the implications for the model when $\zeta$ is "high" (near 1) versus when it is "low" (near 0)? Firstly, it is observed that there is a cutoff $\zeta$ below which the model no longer shows two different states of operation. This is defined as follows:

$$\zeta_{cutoff} = \frac{\varepsilon}{\varepsilon + \nu}$$

Why is this a cutoff value of $\zeta$? This is the minimum value that $\varphi$ can take at any time, since there will always be a spontaneous component present in the input frequency, even when there are not sufficient resources for signaling vesicular release. In other words:

$$\varphi_{min} = \zeta_{cutoff} = \frac{\varepsilon}{\varepsilon + \nu}$$

If $\zeta$ for the network is set lower than $\varphi_{\mathrm{min}}$, then the condition $\varphi > \zeta$ is always true, and the entire network will only show one state of operation (after a transient), which occurs in regimes A and B described below.

The interpretation of the comparison of $\varphi$ to $\zeta$ can be further explored. We again emphasize that the behavior of the presynaptic terminal of interest is representative of the behavior of other synapses of the network. However, every presynaptic terminal in the network will *not* be in exactly the same state at the same time. Rather, they will show a *distribution* of amounts of available ATP and filled vesicles. These distributions need not be normal (Gaussian) in nature, however they will have mean values, and these means can be compared to the values in the representative presynaptic terminal. The quantity $\varphi$ is an indicator of the availability of resources in the representative presynaptic terminal. When $\zeta$ is "high" (near 1), the statement $\varphi > \zeta$ imposes a more stringent condition on the availability of resources in the representative presynaptic terminal. However, since the presynaptic terminal of interest is a representative of all of the presynaptic terminals in the network (which have a distribution of values for available ATP and filled vesicles), the implied condition is that for "high" $\zeta$, the distribution of values for available ATP and filled vesicles cannot be as spread out in order to still fulfill the condition $\varphi > \zeta$—in mathematical terms, the distribution must have lower kurtosis. On the other hand, a "low" $\zeta$ permits $\varphi > \zeta$ to be true at lower values of $\varphi$. This implies that presynaptic terminals with lower available resources still can drive the network in the high frequency state, and the distribution of amounts of available ATP and filled vesicles is allowed to have a higher kurtosis while still meeting the condition $\varphi > \zeta$.

UP and DOWN States

What is the relationship between the critical numbers $\chi$ , $\zeta$, and $\varphi$, and UP and DOWN states in this model? As described above, $\chi$ determines the length of the limit cycle generated within the model. This, then, describes the duration of time that the system will be in one cycle of UP state + DOWN state. As described earlier, $\zeta$ and $\varphi$ (a function of the variable component of the input frequency, $v$) determine which mode (or regime) the system functions in. Thus, the specific combinations of $\zeta$ and $\varphi$ (or $v$) determine whether or not there are even any distinct UP or DOWN states, or whether there is only a single state.

Membrane Potentials

As noted previously, membrane potentials were calculated based on gradients of sodium and potassium ion concentrations across the presynaptic cell membrane, and a DC shift was employed to account for the other ionic species not otherwise modeled. Ion movements occurred due to two processes: 1) synaptic vesicle release and 2) action of the sodium-potassium pump.

The former process was assumed to cause movement of sodium into the presynaptic terminal, and potassium out of the presynaptic terminal, due to the arrival of a virtual action potential (abstracted into the firing frequency). The movement of ions was assumed to be instantaneous—diffusion processes were not modeled. The action of the sodium-potassium pump was modeled using two separate processes, one which extrudes sodium from the presynaptic terminal, and the other which causes uptake of potassium into the presynaptic terminal. The ATP consumption of the sodium potassium pump was also modeled separately as a process that consumes ATP and generates ADP.

UP and DOWN states as defined in the literature have not only the properties of higher firing frequency and lower firing frequency respectively, but also have associated differences in membrane potential. UP states characteristically are associated with a somewhat more depolarized (more positive) resting membrane potential, whereas DOWN states characteristically have a somewhat more hyperpolarized (more negative) resting membrane potential.[13,18] In this model, distinctive UP states (defined by higher firing frequency and more depolarized resting potentials) and DOWN states (defined by lower firing frequency and more hyperpolarized membrane potentials) are generated in regimes C, D, E, and F. Strikingly, the depolarized and hyperpolarized membrane potentials associated with the UP and DOWN states, respectively, are generated in the presynaptic terminal as a ***result*** of the virtual action potentials (modeled by the input firing frequency) causing synaptic vesicle release, rather than the rate of synaptic vesicle release changing ***due to*** alterations in the membrane potential. It has been posited that UP states switch to DOWN states due to a fatigue mechanism such as spike frequency adaptation currents or synaptic short-term depression, and these fatigue mechanisms recover until the network switches back into the UP state.[13] Our model shows that the "fatigue variables" are actually the availability of neurotransmitter and ATP.

The kinetics of specific ion channels has not been modeled, as noted above. Notably, Ching et al.[39] specifically invoked the $K_{ATP}$ channel in their neurophysiologic-metabolic model. This was modeled as an ATP-dependent potassium current using the Hodgkin-Huxley formalism. While this channel was not included in the current model, its potential effect on the presynaptic terminal can be analyzed. Specifically, the $K_{ATP}$ channel opens and causes hyperpolarization of the cell when ATP concentrations decrease. In the context of the model, this would lead to a lower input firing frequency (due to hyperpolarization increasing the threshold for action potential firing; this process is abstracted within the input firing frequency), thereby decreasing ATP use. This, in turn, allows the presynaptic terminal to recover its ATP store more quickly. When the ATP stores recover, the $K_{ATP}$ channel closes, allowing the input firing frequency to rise again (due to the withdrawal of hyperpolarization abstracted away in the input firing frequency). The inclusion of a $K_{ATP}$ channel process will simply augment the homeostatic processes that are already being modeled. *In vivo*, the reality is likely bi-directional: membrane potential changes affect firing rates, and firing rates affect baseline membrane potentials.

Finally, one definition of sleep as noted above[7] is 1) quiescence (decreased activity), 2) increased threshold, and 3) homeostasis. In our model, the membrane potential is a surrogate for increased threshold, because a change in membrane potential does not directly change the rate of synaptic vesicle release. However, one way to capture this effect in the future would be to modulate the input firing frequency as a function of the membrane potential. As has been noted previously, the input firing frequency abstracts all of the processes involved in generating an action potential in the presynaptic cell, including spatial and temporal summation of excitatory and inhibitory inputs. When the balance of excitation to inhibition favors excitation, the input firing frequency increases. When the balance of excitation to inhibition favors inhibition, the input firing frequency decreases. This illustrates the important point that there is no explicit representation of inhibition in our model—it is included in the independent parameter of input firing frequency.

Limit Cycles

The regimes A-F of the firing patterns of the presynaptic terminal will be described in this section. Of note, each simulation run had a "starting transient" which lasted up to approximately 40 seconds, and was included in the figures, but not included in the analysis. Regime A is what we have called "input-limited"—the presynaptic terminal always has both sufficient ATP and a sufficient number of vesicles; the output firing frequency is limited only by incoming input firing frequency. In this situation, the moving average firing frequency perfectly matches the combined maximum input frequency ($\varepsilon + v$), within the precision of the discrete numerical solver. Regime A has been previously illustrated in Figure 2. The membrane potential remains at a single baseline, which is near that of the UP state in other regimes (see below). Notably, our model does not include a limit on the maximum number of vesicles in the presynaptic terminal; this would be determined by volume considerations.

Vesicle-limited cycles

Regime B and Regime C are termed "vesicle limited". This indicates that the presynaptic terminal is able to fire at the same frequency as the input frequency—meaning that the moving average output frequency equals the input frequency—until it runs out of vesicles. In this situation, Regime B and Regime C show different behaviors. In Regime B (Extended Data Figure 6a), ATP remains stable after a transient drop (second trace). The input frequency remains the combined maximum ($\varepsilon + v$) for the entire run (third trace) except for a brief drop to the spontaneous frequency ($\varepsilon$) when the number of vesicles initially drops to one (top trace, note logarithmic scale on *y*-axis). After the initial drop, the number of vesicles continues to oscillate between one and three (top trace). The moving average output frequency (fourth trace) is able to match the input frequency (third trace) while the number of vesicles is still above one (top trace). However, after the number of vesicles drops to one and begins oscillating between one and three, the moving average output frequency (fourth trace) begins oscillating around an intermediate frequency which is neither the combined maximum input frequency ($\varepsilon + \nu$) nor the spontaneous frequency ($\varepsilon$). This is due to missed firing events from lack of filled vesicles; a vesicle is released as soon as it is filled. There is a subtle change in the membrane potential (bottom trace) after the number of vesicles drops to one, and then it remains stable. Extended Data Figure 6b shows a vesicle-limited mode. The limit cycle describes tight loops in the bottom left corner of the graph, as seen previously in Figure 4. In Extended Data Figure 6c, it is seen that plotting the moving average output frequency against the number of filled vesicles yields tight loops that take on an "X" shape due to the vesicle number oscillating between one and two, as the moving average output frequency oscillates between two intermediate values, after a transient.

Regime C is also a "vesicle-limited" mode. In this regime (Extended Data Figure 7a), the number of vesicles oscillates between approximately 90 and zero (top trace). ATP oscillates between a higher and a lower level (second trace). With the exception of a transient at the beginning, it never drops below the ATP threshold for firing (dashed line). The input frequency (third trace) oscillates between the combined maximum frequency ($\varepsilon + v$) and the spontaneous frequency ($\varepsilon$), remaining at the lower spontaneous frequency when the number of vesicles is building up (rising phases of top trace), and attaining the higher combined maximum frequency when the number of vesicles is being depleted (falling phases of top trace). The duration of the limit cycles in this mode, as indicated by the duration of one cycle of vesicle oscillations, is between 30 and 50 seconds. The moving average output frequency (fourth trace) is able to match the input frequency (third trace) at all times except for a brief early transient. The membrane potential (bottom trace) attains two distinct values; the more depolarized UP states correspond to the falling phases in the number of vesicles (top trace) as well as the higher values for input and moving average output frequencies (third and fourth traces). Similarly, the more hyperpolarized DOWN states correspond to the rising phases of the number of vesicles and the lower values for input and moving average output frequencies, when the presynaptic terminal is recovering. Extended Data Figure 7b illustrates the presence of a vesicle-limited mode. The number of vesicles oscillates between approximately 90 and zero, describing large loops. ATP never drops below the ATP threshold for firing (dashed line; seen better in Figure 4). Plotting the moving average output frequency against the number of filled vesicles (Extended Data Figure 7c) yields large well-formed loops as the vesicle number oscillates between one and approximately 90, and the moving average output frequency oscillates between the spontaneous frequency ($\varepsilon$) and the combined maximum input frequency ($\varepsilon + \nu$), after a transient.

ATP-limited cycles

Regime F is a purely “ATP-limited” mode: the presynaptic terminal’s moving average output frequency approximately matches the combined maximum input frequency ($\varepsilon + \nu$) until its ATP is depleted. In Extended Data Figure 10a, the number of vesicles oscillates between approximately 100 and 80, but never reaches zero (top trace). ATP oscillates between a higher and a lower level (second trace), repeatedly dropping below the threshold for firing (dashed line). The input frequency (third trace) rapidly oscillates between the combined maximum frequency ($\varepsilon + \nu$) and the spontaneous frequency ($\varepsilon$), remaining at the lower spontaneous frequency when the

ATP concentration is building up (rising phases of second trace), and attaining the higher combined maximum frequency when the ATP concentration is being depleted (falling phases of second trace). The duration of the limit cycles in this mode, as indicated by the duration of one cycle of ATP oscillations, is much shorter than those seen in regime C—each cycle lasts between 2.5 and 7 seconds—and there a significant variation in duration from cycle to cycle. The moving average output frequency (fourth trace) is only able to briefly match the input frequency (third trace), after which it falls to a low intermediate frequency at approximately 7 Hz, before falling ultimately to the spontaneous input frequency ($\varepsilon$). This is visible in the inset to the fourth trace. The membrane potential (bottom trace) attains two distinct values; the more depolarized UP states correspond to the falling phases in the ATP concentration (second trace) as well as the higher values for input and moving average output frequencies (third and fourth traces). Similarly, the more hyperpolarized DOWN states correspond to the rising phases of the ATP concentration and the lower values for input and moving average output frequencies, when the presynaptic terminal is recovering. Extended Data Figure 10b depicts an ATP-limited mode. The ATP concentration rapidly oscillates between a higher value and the threshold value (dashed line), while the number of vesicles minimally changes, leading to flattened, elongated limit cycles. When the moving average output frequency is plotted against the number of filled vesicles (Extended Data Figure 10c), small trapezoidal loops are formed as the vesicle number oscillates between approximately 80 and 95, and the moving average output frequency oscillates between the spontaneous frequency ($\varepsilon$), the combined maximum input frequency ($\varepsilon + \nu$), and several intermediate frequencies, after a transient.

#### Dual-limited cycles

“Dual-limited” cycles are those which demonstrate features of both limitation due to insufficient ATP and limitation due to insufficient number of vesicles.

##### Vesicle-limited with superimposed ATP limitation

Regime D is a unique mode which demonstrates an ATP limitation superimposed on a vesicle limitation. Extended Data Figure 8a shows that the number of vesicles oscillates between approximately 90 and zero (top trace). ATP oscillates between a higher and a lower baseline level (second trace). At the lower baseline level, the ATP concentration repeatedly drops below the threshold for firing (dashed line). The input frequency (third trace) oscillates between the combined maximum frequency ($\varepsilon + \nu$) and the spontaneous frequency ($\varepsilon$), remaining at the lower spontaneous frequency when the number of vesicles is building up (rising phases of top trace), and attaining the higher combined maximum frequency when the number of vesicles is being depleted (falling phases of top trace). The moving average output frequency (fourth trace) is only able to briefly match the input frequency (third trace), after which it oscillates around a low intermediate frequency at approximately 9 Hz, before falling ultimately to the spontaneous input frequency ($\varepsilon$). The membrane potential (bottom trace) attains two distinct values; the more depolarized UP states correspond to the falling phases in the number of vesicles (top trace) as well as the higher values for input and moving average output frequencies (third and fourth traces). Notably, during the UP states, the ATP concentration oscillates above and below the threshold for firing, indicating a superimposed ATP limitation. This is reflected by the lower moving average output frequencies (fourth trace) during the latter parts of the UP states. The more hyperpolarized DOWN states correspond to the rising phases of the number of vesicles and the lower values for input and moving average output frequencies, when the presynaptic terminal is recovering. The duration of the overall limit cycle, as indicated by the cycles of number of vesicles dropping to zero, is intermediate—23-30 seconds—with a superimposed high frequency oscillation of 0.3-2 seconds due to the ATP limitation. A dual-limited mode is seen in Extended Data Figure 8b. The limit cycles describe large loops, and are difficult to distinguish from those of the vesicle-limited regime. ATP repeatedly drops below the ATP threshold for firing (dashed line). However, plotting the moving average output frequency against the number of filled vesicles (Extended Data Figure 8c) yields large loops clearly distinguishable from those in regime C (Extended Data Figure 7c) as the vesicle number oscillates between one and approximately 90. Unlike in regime C, there is a “step-off” in frequency on the right side of the graph, indicating the intermediate output frequency that results from the superimposed ATP limitation during the UP states. The moving average output frequency oscillates between the spontaneous frequency ($\varepsilon$), the combined maximum input frequency ($\varepsilon + v$), and this intermediate output frequency.

Extended Data Figure 9 shows a more detailed view of one UP state period in regime D. The moving average output frequency (second trace), and the *instantaneous* output frequency (defined as the inverse of the difference of the times of two sequential firing events, third trace) initially both match the input frequency (first trace). They are able to match the input frequency while the ATP concentration is still sufficient (fourth trace). However, after the ATP is depleted to below the threshold level (dashed line), the moving average output frequency (second trace) drops to a lower intermediate level. The instantaneous output frequency gives a more detailed view, where it is seen that the presynaptic terminal can attain a higher intermediate instantaneous frequency (third trace) when ATP is above the threshold (fourth trace), and has a lower instantaneous output frequency when ATP is below threshold. The presynaptic terminal is able to reach the higher instantaneous frequency approximately every second to third firing event (third trace); this is averaged out and leads to a much lower amplitude fluctuation in the moving average output frequency (second trace). When the input frequency (first trace) drops to the low spontaneous frequency ($\varepsilon$), the moving average output frequency (second trace) and instantaneous output frequency (third trace) follow suit, and ATP concentrations recover (fourth trace).

### Transitional dual-limited cycles

Regime E is a transitional mode which demonstrates initial ATP limitation (as in regime F), followed by a dual-limited appearance (as in regime D). Extended Data Figure 11a shows that the number of vesicles initially remains at around 90, but then begins to oscillate between approximately 90 and zero (top trace). Initially ATP rapidly oscillates between a higher and a lower baseline level similar to regime F (second trace). It then switches to a slower oscillation like in regime D. In both segments, at the lower baseline level, the ATP concentration repeatedly drops below the threshold for firing (dashed line). The input frequency (third trace) oscillates between the combined maximum frequency ($\varepsilon + v$) and the spontaneous frequency ($\varepsilon$). Initially it oscillates between levels in sync with ATP concentration (approximately first 40s of second trace), but then it starts oscillating at the same rate as the number of vesicles (top trace, after ~40s). The moving average output frequency (fourth trace) is only able to briefly match the input frequency both before and after ~40s (third trace), after which it oscillates around a low intermediate frequency, before falling ultimately to the spontaneous input frequency ($\varepsilon$). The membrane potential (bottom trace) attains two distinct values; it rapidly oscillates between more depolarized UP states and more hyperpolarized DOWN states when the system is limited by ATP (approximately first 40s, as seen in second trace), and then oscillates more slowly when the system is dual-limited (after approximately 40s, as seen in top and second traces). Extended Data Figure 11b reveals a transitional mode. The limit cycles initially start as short elongated regime F-like loops (blue arrow), and then transition to large regime D-like loops (black arrow).

Similarly, in Extended Data Figure 11c, plotting the moving average output frequency against the number of filled vesicles yields large loops, the first of which appear F-like (blue arrow), the latter of which are D-like (black arrow). The moving average output frequency oscillates between the spontaneous frequency ($\varepsilon$), the combined maximum input frequency ($\varepsilon + v$), and an intermediate output frequency, which arises due to superimposed ATP limitation.

There were several other transitional modes which were less frequently seen, at the borders between the other established modes (regimes B, C, D, and F). These included modes where the behavior started as in regime C, but then transitioned to regime B ("BC"), and modes where the behavior started as in regime D, but then transitioned to regime B ("BD") (*data not shown*). A rarer transitional mode was where the behavior started in the first transitional mode E (where the behavior starts as in regime F, but then switches to regime D), but then switched back to behavior as in regime F (*data not shown*). In many of these other transitional modes, the moving average output frequency only briefly reaches the input frequency ($\varepsilon + v$), and then falls to a lower intermediate value which is significantly higher than the low spontaneous rate ($\varepsilon$), during the UP state (*data not shown*).

Duration of Limit Cycles

The durations of the UP and DOWN states can now be examined (Extended Data Figure 14). Neither Regime A nor Regime B have discrete UP or DOWN states. For regimes C-F, the DOWN state duration depends on availability of glutamate and ATP. Neither is rate-limiting through regimes C-E, so the DOWN state duration minimally changes as a function of input frequency. In regime F, the duration of the DOWN state depends more on ATP availability, and decreases because ATP cycling occurs more rapidly than cycling of filled vesicles.

Regime B does not have a discrete UP state, but it does have an initial state where the moving average output frequency is equal to the input frequency. The duration of this single "high frequency" state, which represents the starting transient for this regime, drops inversely with frequency and also depends on the initial conditions of the model. This dependence on initial conditions is the reason for the discontinuity between the durations for regimes B and C. For regimes C-F, the limit cycle durations are calculated based on the stationary state (excluding the initial transient) of the model. The UP state duration is strongly a function of input frequency in regime C (Extended Data Figure 14), because it depends on how rapidly the vesicle pool is depleted. There is a slight nonlinearity because vesicles continue to fill during the UP state, at a lower rate. The duration of the UP state is essentially inversely proportional to the input frequency. In regimes D and E, ATP limitation caps the effective output frequency (moving average frequency of synaptic vesicle release) to a near-constant, which limits the duration of the UP state to a near-constant. The UP state drops significantly in duration in regime F, due to the stronger dependence on ATP availability rather than vesicle availability.

Our model qualitatively recapitulates results from both an *in vitro* model and a computational model based on the Hodgkin-Huxley formalism. Chen *et al* showed that in a cortical slab model, decreasing inhibition by adding the $GABA_A$ antagonist bicuculline, increased the frequency of active states, and significantly shorted the duration of each active state. In their computational model, they demonstrated that by decreasing the synaptic strength of feedback inhibition, initially there was an increase in active state duration. This was followed by a decrease in active state duration with intense bursting, with further increased inhibition.[43] In both cases, the decrease of inhibition corresponds with an ***increase*** in the variable portion of the input frequency (*v*). As in Chen's work, our model shows that there is initially an appearance of UP and DOWN states with increasing input frequency. Unlike their initial finding of an increasing duration of UP states, our model shows that this duration is inversely proportional to input frequency—this may be explained by a neuronal "reserve" in the *in vitro* system, the kinetics of the antagonist used, or may be due to the method of recording. However, after the input frequency increases sufficiently, the model transitions into a mode where the duration of the UP state significantly decreases (regime F), just as seen in both the *in vitro* and *in silico* models by Chen *et al*. This suggests that the intense bursting and shortening of UP state duration in the models by Chen *et al* is due to their systems becoming ATP-limited due to loss of inhibition.

### Implications for Organism Level Sleep

The existence of multiple modes of operation begs the question: "Which regime do neurons operate in *in vivo*?" This is a difficult question to answer, and it is likely that neurons in different parts of the brain operate in different regimes at different times, depending on the computational demands of that region of the brain. An important factor to consider is that our model utilizes a constant source of glucose, with an ability to produce any amount of both ATP and glutamate (limited only by the reaction rates for the synthesis of the two species). This means that the regimes described above can recur indefinitely. This indefinite progression is certainly non-physical, as glucose and amino acid supplies are limited *in vivo*. What likely occurs is that neurons function in one of the above regimes until they can no longer be supplied sufficient raw resources (i.e. glutamine, glucose, and other nutrients necessary for function), then they drop into a further low state of function until they can recover those resources. This adds another layer of use-dependence, which has been seen *in vivo* (for example, see Vyzazovskiy *et al*).[29] One intriguing possibility is that neurons function in one of the regimes that allow brief bursts of extremely high frequency activity (regime D, E, or F), as an adaptive mechanism for organisms to quickly respond to rapidly changing stimuli in the environment (such as, say, the appearance of a predator). Since this sort of stimulus that needs to be rapidly processed is relatively uncommon, the organism can “afford” to use up all of its neuronal resources briefly in order to survive the situation, and recovers those resources by local and global (behavioral) sleep.

### Role of Diffuse Modulatory Systems

Our model focuses exclusively on a glutamatergic synapse, and can be easily extended to represent a GABAergic synapse. These two neurotransmitters together comprise approximately 90% of the neurons in the brain.[5] However, numerous studies have shown the importance of what have been termed the “diffuse modulatory systems” in the generation and maintenance of sleep at the organism level; some of these studies will be reviewed below. The diffuse modulatory systems involved in sleep include serotonergic, histaminergic, cholinergic, noradrenergic, and orexinergic systems.

### Circadian Drive versus Homeostatic Drive

A two-process model of sleep regulation was originally proposed many years ago to explain the behavior of the sleep cycle in animals. The two processes, a homeostatic Process S and a circadian Process C, interact together to determine when and for how long an organism sleeps. Process S is conceptualized as “sleep debt”, which increases during wakefulness and decreases during sleep, and Process C serves to entrain the cycle to day and night. The original formulation does not include observed phenomena such as regional use-dependent homeostasis or the presumed mutual interactions between Process S and Process C, but it remains a helpful construct.[44] Other models with homeostatic mechanisms have been proposed as well, including extension of the original two-process model (such as the three-process model), reciprocal interaction models, limit cycle reciprocal interaction models, and combinations.[45] It is noted that all of these models attempt to explain sleep cycling at a network level, such as describing the role of the suprachiasmatic nucleus in circadian rhythms, without reference to the cellular level. Nevertheless, even at the network level, some groups have connected the modulation of rest- and activity-time with changes in metabolism.[44] Indeed, in one human study it was found that the amount of activity during the wakeful period remained roughly stable regardless of the *duration* of the wakefulness, and it was suggested that the basal metabolic rate varies inversely with the duration of wakefulness.[46]

Flip-flop Circuit for Sleep

Another major model has been advanced, which falls under the rubric of a reciprocal interaction model, or limit cycle reciprocal interaction model, as listed above. This is the concept of a "flip flop" circuit for sleep, wherein there are mutually inhibitory sleep circuits that allow the brain to transition between states. Wake-promoting networks are thought to consist of cholinergic, monoaminergic (noradrenergic, serotoninergic, and dopaminergic), and histaminergic neurons. Another important neurotransmitter in wakefulness is orexin; orexinergic neurons are also driven by low glucose levels, and may be involved in generating foraging behaviors in awake animals. Specific glutamatergic neurons are also thought to be involved in promoting wakefulness.[47] NREM-active neurons include those in the ventrolateral preoptic nucleus (VLPO). There are GABAergic and galaninergic neurons in this region, but it has been difficult to determine specific cellular identities in single-cell recordings. These neurons receive reciprocal input from the neurons residing in various wake-promoting regions. Finally, certain neurons, such as cholinergic neurons, appear to be most active during REM sleep. The connection patterns of these neurons have been proposed to form the “flip flop switch” between wake-promoting and sleep-promoting neurons, similar to REM-on and REM-off neurons.[47] Interestingly, multiple factors have been suggested to cause the “flip flop switch” to flip, such as adenosine as a marker for ongoing cellular metabolism[48], and orexin as a marker for hypoglycemia and/or a fasted state.[49]

### Do Diffuse Modulatory Systems also have Rest-Activity Cycles?

As noted above, in both the two-process model and the “flip flop switch” models, there is a potential for metabolic factors to affect the state of the brain. However, this remains at the network level—the level at which neurons using different neurotransmitters interact with one another. Our model addresses the necessity of glutamatergic neurons to “sleep” (have rest and activity periods), and the metabolic processes will be relatively similar in GABAergic neurons. What about neurons in the diffuse modulatory systems? Do they also have periods of rest and activity due to resource limitation? We are not aware of any study that has directly investigated this question. However, it has been noted that of the different neurons thought to be participating in sleep cycling, specific noradrenergic, dopaminergic, serotoninergic, and histaminergic neurons generally fire most during wakefulness, less during NREM sleep, and almost not at all during REM sleep. Orexinergic neurons are also more active during wakefulness, and drive the activity of many of the previously mentioned neurons. On the other hand, neurons in the VLPO are primarily active only during sleep.[50] This indicates that, at the very least, the neurons of the diffuse modulatory systems involved in sleep have the opportunity to go into a low-activity mode wherein they can restore their energy supply and vesicle pools.

References:


24 Tramm, N., Oppenheimer, N., Nagy, S., Efrati, E. & Biron, D. Why do sleeping nematodes adopt a hockey-stick-like posture? *PloS one* **9**, e101162, doi:10.1371/journal.pone.0101162 (2014).

25 Krueger, J. M. *et al.* Sleep as a fundamental property of neuronal assemblies. *Nature reviews. Neuroscience* **9**, 910-919, doi:10.1038/nrn2521 (2008).

26 Cirelli, C. & Tononi, G. Is sleep essential? *PLoS biology* **6**, e216, doi:10.1371/journal.pbio.0060216 (2008).

27 Lyamin, O. I., Kosenko, P. O., Lapierre, J. L., Mukhametov, L. M. & Siegel, J. M. Fur seals display a strong drive for bilateral slow-wave sleep while on land. *The Journal of neuroscience : the official journal of the Society for Neuroscience* **28**, 12614-12621, doi:10.1523/JNEUROSCI.2306-08.2008 (2008).

28 Rattenborg, N. C. *et al.* Evidence that birds sleep in mid-flight. *Nature communications* **7**, 12468, doi:10.1038/ncomms12468 (2016).

29 Vyazovskiy, V. V. *et al.* Local sleep in awake rats. *Nature* **472**, 443-447, doi:10.1038/nature10009 (2011).

30 Nichols, A. L. A., Eichler, T., Latham, R. & Zimmer, M. A global brain state underlies C. elegans sleep behavior. *Science* **356**, doi:10.1126/science.aam6851 (2017).

31 Nath, R. D. *et al.* The Jellyfish Cassiopea Exhibits a Sleep-like State. *Current biology : CB* **27**, 2984-2990 e2983, doi:10.1016/j.cub.2017.08.014 (2017).

32 Field, J. M. & Bonsall, M. B. The evolution of sleep is inevitable in a periodic world. *PloS one* **13**, e0201615, doi:10.1371/journal.pone.0201615 (2018).

33 Krishnan, G. P. *et al.* Cellular and neurochemical basis of sleep stages in the thalamocortical network. *eLife* **5**, doi:10.7554/eLife.18607 (2016).

34 Krueger, J. M., Nguyen, J. T., Dykstra-Aiello, C. J. & Taishi, P. Local sleep. *Sleep medicine reviews* **43**, 14-21, doi:10.1016/j.smrv.2018.10.001 (2019).

35 Halasz, P., Terzano, M., Parrino, L. & Bodizs, R. The nature of arousal in sleep. *Journal of sleep research* **13**, 1-23 (2004).

36 Patel, A. B. *et al.* Glutamatergic neurotransmission and neuronal glucose oxidation are coupled during intense neuronal activation. *Journal of cerebral blood flow and metabolism : official journal of the International Society of Cerebral Blood Flow and Metabolism* **24**, 972-985, doi:10.1097/01.WCB.0000126234.16188.71 (2004).

37 Tewari, S. G. & Majumdar, K. K. A mathematical model of the tripartite synapse: astrocyte-induced synaptic plasticity. *Journal of biological physics* **38**, 465-496, doi:10.1007/s10867-012-9267-7 (2012).

38 Flanagan, B., McDaid, L., Wade, J., Wong-Lin, K. & Harkin, J. A computational study of astrocytic glutamate influence on post-synaptic neuronal excitability. *PLoS computational biology* **14**, e1006040, doi:10.1371/journal.pcbi.1006040 (2018).

39 Ching, S., Purdon, P. L., Vijayan, S., Kopell, N. J. & Brown, E. N. A neurophysiological-metabolic model for burst suppression. *Proceedings of the National Academy of Sciences of the United States of America* **109**, 3095-3100, doi:10.1073/pnas.1121461109 (2012).

40 Hodgkin, A. L. & Huxley, A. F. A quantitative description of membrane current and its application to conduction and excitation in nerve. *The Journal of physiology* **117**, 500-544, doi:10.1113/jphysiol.1952.sp004764 (1952).

41 Steriade, M., Timofeev, I. & Grenier, F. Natural waking and sleep states: a view from inside neocortical neurons. *Journal of neurophysiology* **85**, 1969-1985, doi:10.1152/jn.2001.85.5.1969 (2001).

42 Hennig, M. H. Theoretical models of synaptic short term plasticity. *Front Comput Neurosci* **7**, doi:10.3389/fncom.2013.00154 (2013).

43 Chen, J. Y., Chauvette, S., Skorheim, S., Timofeev, I. & Bazhenov, M. Interneuron-mediated inhibition synchronizes neuronal activity during slow oscillation. *The Journal of physiology* **590**, 3987-4010, doi:10.1113/jphysiol.2012.227462 (2012).

44 Borbely, A. A., Daan, S., Wirz-Justice, A. & Deboer, T. The two-process model of sleep regulation: a reappraisal. *Journal of sleep research* **25**, 131-143, doi:10.1111/jsr.12371 (2016).

45 Achermann, P. & Borbely, A. A. Mathematical models of sleep regulation. *Frontiers in bioscience : a journal and virtual library* **8**, s683-693, doi:10.2741/1064 (2003).

46 Aschoff, J. On the relationship between motor activity and the sleep-wake cycle in humans during temporal isolation. *Journal of biological rhythms* **8**, 33-46, doi:10.1177/074873049300800103 (1993).

47 Saper, C. B., Fuller, P. M., Pedersen, N. P., Lu, J. & Scammell, T. E. Sleep state switching. *Neuron* **68**, 1023-1042, doi:10.1016/j.neuron.2010.11.032 (2010).

48 Porkka-Heiskanen, T. *et al.* Adenosine: a mediator of the sleep-inducing effects of prolonged wakefulness. *Science* **276**, 1265-1268, doi:10.1126/science.276.5316.1265 (1997).

49 Yamanaka, A. *et al.* Hypothalamic orexin neurons regulate arousal according to energy balance in mice. *Neuron* **38**, 701-713, doi:10.1016/s0896-6273(03)00331-3 (2003).

50 Fuller, P. M., Gooley, J. J. & Saper, C. B. Neurobiology of the sleep-wake cycle: sleep architecture, circadian regulation, and regulatory feedback. *Journal of biological rhythms* **21**, 482-493, doi:10.1177/0748730406294627 (2006).
